# Interactions between temperature and nutrients determine the population dynamics of primary producers

**DOI:** 10.1101/2023.08.22.554290

**Authors:** Carling Bieg, David Vasseur

**Affiliations:** Department of Ecology and Evolutionary Biology, Yale University

**Keywords:** Thermal performance, carrying capacity, population dynamics, phytoplankton, Droop model, nutrient limitation, theoretical ecology

## Abstract

Global change is rapidly and fundamentally altering many of the processes regulating the flux of energy throughout ecosystems and although researchers now understand the effect of temperature on key rates (such as aquatic primary productivity), the theoretical foundation needed to generate forecasts of biomass dynamics and extinction risk remains underdeveloped. We develop new theory that describes the interconnected effects of nutrients and temperature on phytoplankton populations and show that the thermal response of equilibrium biomass (i.e., carrying capacity) always peaks at a lower temperature than for productivity (i.e., growth rate). This difference results from trade-offs between the thermal responses of growth, death, and per-capita impact on the nutrient pool, making this thermal mismatch highly general and applicable to widely used population models. We further show that non-equilibrium dynamics depend on the pace of environmental change relative to underlying vital rates, and that populations respond to variable environments differently at high vs. low temperatures due to thermal asymmetries.

## Introduction

Global change is rapidly altering the abiotic environment in multiple ways, including changes in the mean and variability of temperatures and suites of other anthropogenic impacts like eutrophication (Doney *et al*. 2012; Steffen *et al*. 2015). Within the ecological hierarchy, populations lie at the interface between abiotic environmental changes and biotic community or ecosystem dynamics, integrating and coupling various responses to environmental change. Despite this, we lack a mechanistic understanding of how population dynamics and resilience respond to multiple axes of global change (i.e., changes in the environment across multiple niche axes) and especially to combinations of abiotic and biotic stressors. The multitude of rapidly changing environmental conditions is effectively altering multiple niche axes at once, creating novel environments and highlighting the importance of understanding the interaction between multiple “stressors.” Critically, our current theory is not equipped to understand population structure and persistence under multiple simultaneous environmental changes, making it difficult to forecast population dynamics in today’s changing, and increasingly variable, world. For example, while the thermal dependence of vital rates has long been acknowledged as important in regulating population performance or fitness (i.e., growth), and researchers have begun understanding the interactive nature of multiple stressors for some population-level processes, it remains to be seen how populations will dynamically respond to suites of anthropogenic impacts associated with global change. Similarly, it remains to be seen how environmental variability along these niche axes will alter population responses. Indeed, environmental variability can have drastically different effects on population performance than what would be predicted in static environments (Vasseur *et al*. 2014; Bernhardt *et al*. 2018b; Slein *et al*. 2023).

Researchers are beginning to come to the consensus that there is an interactive effect of temperature and resource limitation on population performance. Specifically, a species’ optimal temperature for growth, as well as the critical limits of its thermal niche, are functions of resource availability such that populations are more sensitive to increasing temperature when resources are limited (Thomas *et al*. 2017; Bestion *et al*. 2018; Huey & Kingsolver 2019; Vinton & Vasseur 2022). While clearly important for intrinsic rates of population growth, it is less clear how this interaction translates to population-dynamic processes such as the thermal response of biomass and long-term population persistence. That is, while thermal performance curves relate directly to population rates of change under idealized (i.e., density-independent) conditions, additional information on density- or resource-dependent population growth is needed to determine the size, dynamics, and extinction risk of populations. Importantly, understanding trends in population biomass is central to population forecasting and necessary for management at various scales. For example, population decline is the key element used to determine a species’ extinction risk (e.g., IUCN status), and biomass is central to dictating the flow of energy throughout food webs and whole ecosystems (i.e., energy flux and carbon or nutrient cycling via numerical responses). Developing theory to understand populations’ thermal and multi-stressor responses at the scale of management is critical for forecasting – and mitigating – the effects of global change.

Empirically, the thermal dependence of population equilibrium biomass (i.e., carrying capacity) is somewhat ambiguous, with evidence ranging from invariant (Jarvis *et al*. 2016) to negative (Fussmann *et al*. 2014; Bernhardt *et al*. 2018a) relationships, while theoretically a variety of nonlinear relationships have been suggested (Savage *et al*. 2004; Amarasekare 2015; Uszko *et al*. 2017; Lemoine 2019; Vasseur 2020). Vasseur (2020) set the logical context for why carrying capacity ought to follow a unimodal relationship with temperature given density- and temperature-dependent birth and death rates, but empirical evidence for this is still lacking as few researchers aim to measure carrying capacity near thermal extremes (and with high enough resolution to capture the thermal limits) – perhaps in large part because of the inherent experimental challenges in doing so – as well as variability in how carrying capacity is measured empirically. More recently, (Vinton & Vasseur 2022) described the thermal dependence of long-term (equilibrium) behaviour of populations under limiting resources, confirming this unimodal relationship and showing that carrying capacity is dependent on both temperature and resource availability. Despite this new understanding of how long-term behaviour ought to respond to multiple stressors, population dynamics involve nonlinearities that make equilibrium behaviour just the starting point for forecasting responses to variable environments (Hastings *et al*. 2018).

Fundamentally, population patterns depend on a dynamic integration of both past and current population densities, abiotic conditions, and trade-offs between costs (metabolism, death) and benefits (somatic and reproductive growth) of functioning within a given environment. These interacting factors regulate population rates of change (i.e., the speed and direction of changes in biomass) and generate inherently lagged responses to changing environments. Under the rapid pace of global change, populations may or may not be able to “keep up” with changing environmental conditions, while also responding to the underlying variability that characterizes natural environments (i.e., the various frequencies of environmental noise; Dillon *et al*. 2016). Some organisms may be able to adapt or acclimate to different environments, while others are less equipped to do so. The importance of environmental acclimation in regulating populations’ performance has received more attention recently (e.g., (Fey *et al*. 2021; Layden *et al*. 2022)), however mechanistic yet generalizable insight into how this impacts population dynamics remains to be seen. Ultimately, acclimation potential dictates an organism or population’s ability to persist in changing environments, but physiological trade-offs determine long-term implications for persistence. Recently, Anderson et al. (unpublished) showed that thermal acclimation of phytoplankton growth can be explained as a dynamic interplay between temperature, nutrient availability and nutrient storage, giving important mechanistic insight into how populations may respond to changing environments. It remains to be seen how these interacting dynamics and environmental legacy effects translate to longer-term population dynamics – that is, population rates of change as well as biomass trajectories and extinction risk. Effective population forecasting in a time of global change requires an integration of mechanistic organismal research and efficient generalizable population dynamics theory.

In this paper, we begin to do so by integrating recent empirical insights on the thermal dependence of various vital rates within a generalizable framework for population dynamics of phytoplankton populations under limiting nutrients. As the base of all aquatic food webs and a vital element of global carbon cycles, phytoplankton have critical functional importance and have become the hallmark for studying both metabolic/thermal ecology and for experimentally testing theoretical predictions, making them an excellent starting point for developing a mechanistically-informed general theory. Here we use a nutrient- and temperature-dependent Droop model (Droop 1974, 1977; Sauterey & Ward 2022), recently amended and empirically-validated by Anderson et al. (unpublished), to explore how temperature and nutrient limitation collectively impact populations in both constant and variable thermal environments. This model is useful because it is well equipped to understand limiting factors for population growth and biomass accrual, as it separates the rates of nutrient uptake and assimilation via a dynamic cellular nutrient quota – both rates that are now known to be differentially-regulated by temperature. Anderson et al. (unpublished) showed that by differentiating uptake and assimilation processes, temperature limitation can occur at different steps creating thermally-dependent bottlenecks for population growth and suggesting the potential for complex population dynamics.

Importantly, our work relates to understanding nonequilibrium (e.g., transient, seasonal) dynamics, in lakes and marine systems where primary productivity is important for the food web, for carbon storage and carbon sequestration, as well as better understanding harmful algal blooms (HABs), which are becoming increasingly prevalent and seemingly connected to nutrients and temperature. This research provides mechanistic insight into phytoplankton population dynamics under global change, with implications for whole ecosystem functioning. Simultaneously, we use a combination of analytical and numerical approaches, allowing us to make generalizable conclusions consistent with phenomenological modeling approaches. As such, our insights also motivate the inclusion of more realistic environmental context into our general population models.

## The Model

To explore the interaction between nutrient availability and temperature on population growth, biomass and dynamics, we use a 3-dimensional system of ordinary differential equations to describe the coupled dynamics of nutrient (N) availability, intracellular nutrient flux modeled using a dynamic quota (Q) and population biomass of phytoplankton (B). We build off the framework first described by Droop (1974; 1977), which was recently amended by Anderson et al. (unpublished) to incorporate the role of temperature (T) (Figure 1). The dynamic nutrient quota component in this model separates nutrient uptake and assimilation into two separate, temperature-dependent processes (Figure 1). This is useful because it more accurately accounts for nutrient-phytoplankton interactions when not at a steady state (Droop 1977), allowing us to explore non-equilibrium dynamics while retaining the analytical tractability of equilibrium solutions of more simple models (Cunningham & Nisbet 1980; Grover 1992; Smith & Waltman 1994). In this model, nutrient drawdown and biomass accrual is largely regulated by nutrient accessibility – that is, the availability and uptake rate of nutrients.

**Figure 1.**
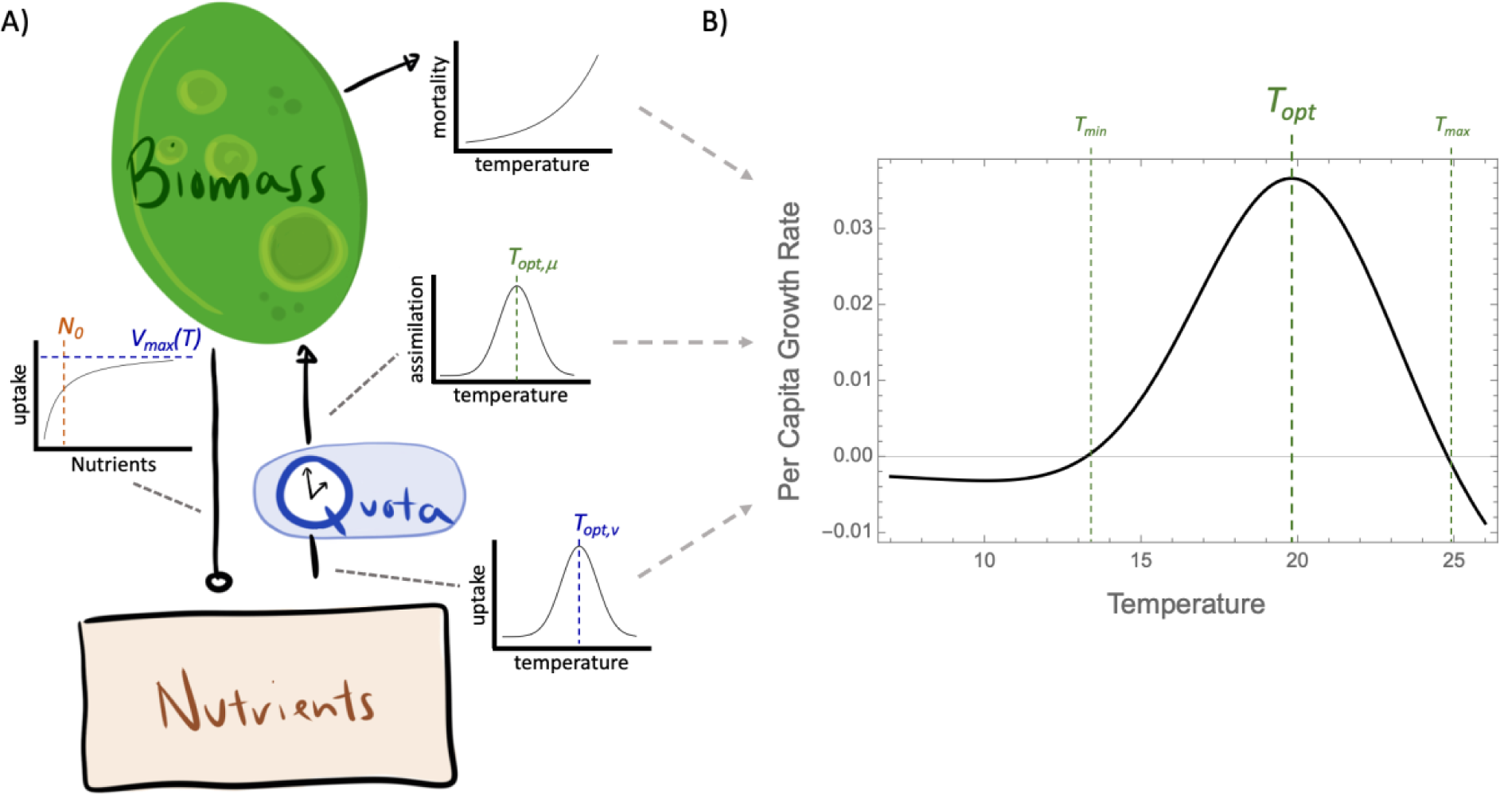
A) Schematic showing model state variables and temperature dependence of various rates. Here, the inclusion of the nutrient quota splits uptake and assimilation into two separate temperature-(and density-) dependent processes, and therefore acts to create a lag in the conversion of nutrients into biomass. The upward arrow indicates conversion of nutrients into biomass (vial the intracellular nutrient quota), and the maximum rate of nutrient uptake (negative interaction, open circle) follows a saturating function of nutrient concentration. B) Collectively, these temperature-dependent rates define a population’s thermal performance curve (TPC), which is defined as the per-capita rate of growth at near-zero densities (i.e., non-limiting nutrients).

Nutrient assimilation, which determines the rate at which stored nutrients (via Q) are converted into biomass, regulates the magnitude of quota build-up, and together uptake and assimilation rates (both of which are temperature dependent) create a trade-off determining the accumulation (rate and magnitude) of an internal nutrient pool via the quota. More broadly, the quota regulates the total flux from resource (nutrients) to biomass and is determined by the balance between density-(and temperature-) dependent uptake and assimilation rates, as well as the available external nutrient pool. While the dynamic nutrient quota has no explicit loss term included here, growth is now scaled by an internal nutrient pool relative to some minimum level required for positive assimilation, and in turn this term can create a lag in population biomass responses to changing environmental conditions.

### Model Equations

Here, nutrient availability (N) is modeled as a chemostat, with nutrient uptake by phytoplankton (B) following a type-II functional response (Monod function), thus allowing for saturation in per-capita uptake:

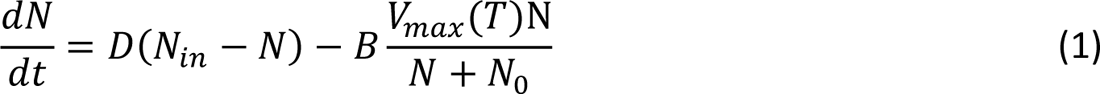

where, *N_0_* is the half-saturation constant for nutrient uptake, *D* is the dilution rate, and *N_in_* is the concentration of nutrients entering the system. *V*_*max*_(*T*) is the temperature-dependent maximum rate of nutrient uptake (moles of nutrient per unit of algal biomass per unit of time. Previous work has shown that *V_max_* (or, more broadly, consumption when not distinguished from assimilation) is a unimodal function of temperature (see (Englund *et al*. 2011) for meta-analysis) and following the work of (Amarasekare & Savage 2012; Thomas *et al*. 2017; Huey & Kingsolver 2019; Vinton & Vasseur 2022), we define *V_max_* as a normally-distributed function of temperature around some optimal temperature for nutrient uptake, *T_opt,v_*:

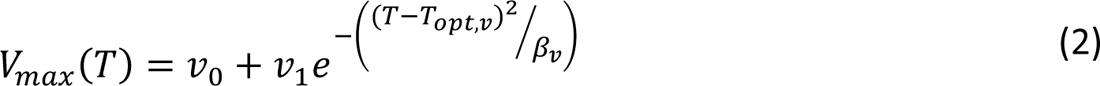

where *β_v_* defines the breadth of the temperature-response for uptake.

Within the cell, the nutrient quota, Q, determines the flux of nutrients based on differences between temperature-dependent uptake and assimilation rates:

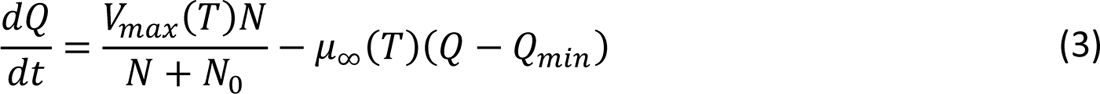

where *Q*_*min*_ is the minimum nutrient quota (moles cell^-1^) needed to maintain a positive assimilation rate and μ_∞_(*T*) is the temperature-dependent maximum rate of nutrient assimilation (time^-1^), defined again as a normally-distributed function of temperature around some optimal temperature for assimilation (*T_opt,μ_*):

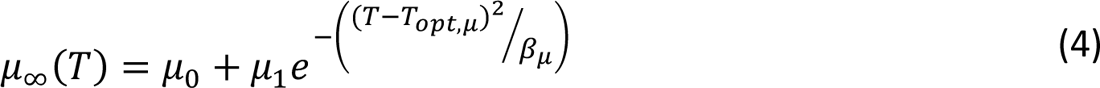

*β_μ_* defines the breadth of the temperature-response for assimilation. μ_∞_ is so-named because it represents the rate of per-capita biomass growth that is achieved when the nutrient quota is infinitely large. Biomass dynamics are thus described as:

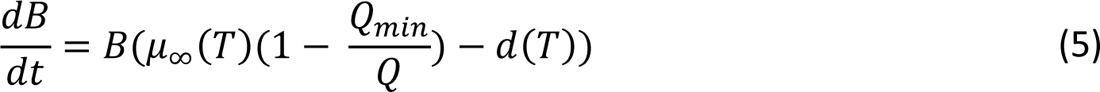

where B is population biomass density (volume^-1^; interchangeable with cell density since cell size is not considered in this model), and *d*(*T*) is the temperature-dependent mortality rate (time^-1^). Previous work has shown that mortality rates scale as Boltzmann-Arrhenius relationships (Gillooly *et al*. 2001; Brown *et al*. 2004; McCoy & Gillooly 2008); however, similar to other theoretical work (Amarasekare 2015; Vinton & Vasseur 2022) we represent mortality as an exponential function of temperature to increase model tractability without losing much accuracy over biologically relevant temperature ranges:

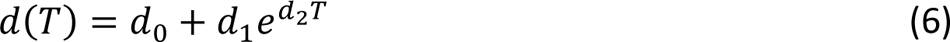

Together, these temperature-dependent rates define the population’s thermal performance.

Although we represent the temperature-dependent functions *V_max_*(*T*) and μ_∞_(*T*) as symmetric unimodal functions, others have assumed that these are monotonically increasing functions of temperature (e.g. Boltzmann-Arrhenius functions; (Thomas *et al*. 2017)). We show in the Appendix that either choice of function leads to a similarly shaped thermal performance curve for phytoplankton growth and that the qualitative patterns demonstrated by our model are not affected by our choice of a unimodal function (Figure A3).

### Numerical & Analytical Analyses

Importantly, our analysis and conclusions are generalizable, particularly due to the analytical approaches we use. That is, while the parameters used in our analysis are loosely based on empirical measurements of important rates, the qualitative relationships between variables gives us insight into phytoplankton growth and population dynamics in general.

When not explicitly stated (e.g., if individually varied for analytical/simulation purposes) the following parameter set was used: *T_opt,v_* = 20, *T_opt,μ_* = 20, σ_v_ = 3.25, σ_μ_ = 3.25, *v_0_* = 0.0005, *v_1_* = 0.005, *μ*_0_ = 0.1, *μ*_1_ = 0.5, *d_0_* = 0.005, *d_1_* = 0.0012, *d_2_* = 0.1, *Q_min_* = 0.1, *r_0_* = 0.5, *N_in_* = 1 and *D* = 1. By setting *D* and *N_in_* to 1, we normalize the inputs of the chemostat model and reduce its dimensionality. We then explore the role of nutrient limitation by varying the half-saturation constant, *N_0_* (uptake saturation), relative to the normalized parameters (as *N_0_* increases for a given nutrient concentration, N, growth is more limited by nutrients). Although uptake saturation and *V_max_* often vary in concert (Aksnes & Egge 1991), we keep these terms separate to better isolate the effects of temperature-dependent uptake and uptake efficiency (1/*N_0_*) as a tractable way of imposing nutrient limitation. Note that in certain applications of the chemostat model, *D* is often incorporated into the population’s mortality or loss term to reflect individuals being washed out; however, consistent with other theory studying biomass dynamics (León & Tumpson 1975; Sauterey & Ward 2022; Vinton & Vasseur 2022) and so that we can better isolate the effects of temperature on dynamics, we assume that phytoplankton mortality is independent of the flow rate *D*. The inclusion of additional mortality to reflect wash-out would not alter general patterns on population responses (i.e., qualitative responses to changing environments) so long as we also include an additive temperature-dependent effect on mortality. The chosen parameter set allows for population persistence (positive growth; stable interior equilibrium) across a reasonable thermal breadth, as shown in Figure 1B, and allows us to qualitatively explore the effects of varying multiple parameters on population persistence and dynamics relative to this baseline.

While the individual rates used in this model are scaled by temperature and/or nutrient availability, we note that this format has not qualitatively changed the set of possible outcomes of the model (i.e., possibility of and qualitative stability properties of stable interior equilibrium). That is, since the model structure has not changed, the possible outcomes of our model are restricted to a set of well-understood phenomena (Droop 1977; Smith & Waltman 1994); Anderson et al. unpublished). Equilibrium solutions for all state variables are described in the Appendix (Equations A1-3), along with the isoclines depicting how temperature changes the general equilibrium structure (Figure A1). This model has two possible stable equilibrium states, depending on the availability of and ability to take up nutrients, one with algae absent (B=0) and one with a positive biomass (B>0) (Figure 2) (Droop 1977; Cunningham & Nisbet 1980; Nisbet & Gurney 1982). In the case where B>0 the population always approaches the equilibrium monotonically and the model does not produce any complex (i.e., cyclic) dynamics. This equilibrium structure therefore reflects the asymptotic behaviour of all simulations under static environmental conditions, and simultaneously serves as a reference for simulation results when environmental variability is incorporated. That is, we refer to all simulation results relative to underlying analytical solutions (i.e., the deterministic skeleton; (Higgins *et al*. 1997)) to further emphasize the generality of our approach and results. Here, rather than focusing on the effects of specific parameterizations (e.g., using an empirically parameterized model or conducting a full parameter sensitivity analysis) we instead focus on the general nature of temperature and nutrient interactions.

**Figure 2.**
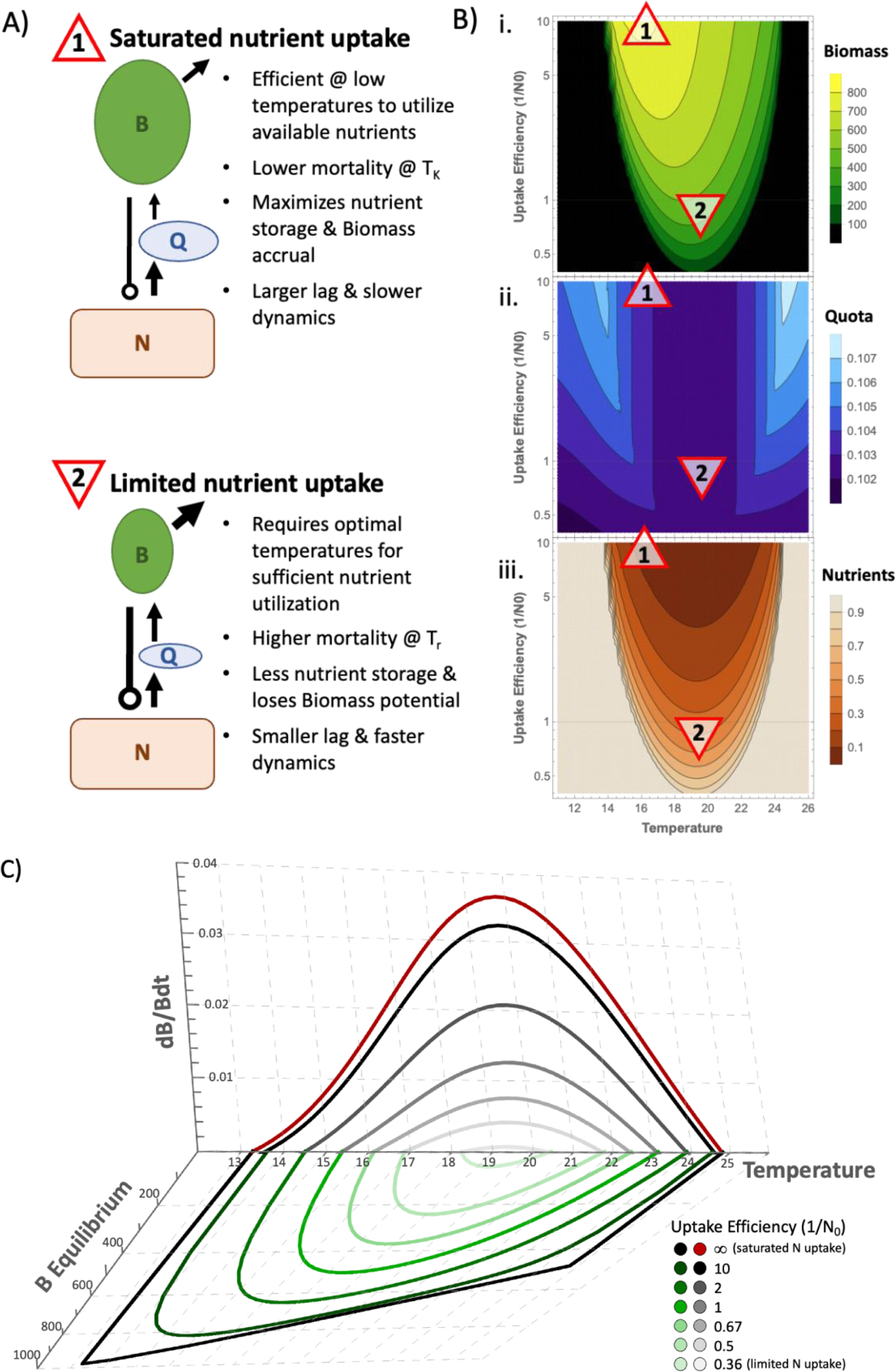
A) Schematic showing the combined influence of nutrient uptake efficiency (defined as 1/N_0_, therefore describing the saturation of uptake) and temperature and their interactive effect in regulating the various rates defining our system; and B) corresponding Nutrient, Quota and Biomass equilibria. C) Temperature responses of productivity (i.e., the TPC; red line representing the fundamental TPC – i.e., infinite nutrients – and black-grey lines representing productivity (i.e., the realized TPC) under changing uptake efficiencies when N=N_in_=1) and equilibrium biomass (green lines), at different levels of nutrient uptake efficiency, defined as 1/N_0_. Specifically notice the temperature mismatch between the two temperature optima, T_r_ and T_K_, for the TPC and equilibrium biomass (i.e., carrying capacity; K), respectively. Here, N_0_ is varied with line/point opacity reflecting limitation of nutrient uptake (higher uptake efficiency, 1/N_0_, effectively equates to saturating nutrients). From light to dark colouring: N_0_ = 0.1, 0.5, 1.0, 1.5, 2.0.

All analyses were done in Wolfram Mathematica v13.1. Numerical simulations were performed using Mathematica’s NDSolve function with automatic integration method. Simulations were run for sufficient time to account for any transient dynamics before evaluating asymptotic behaviour. This duration depended on the analysis being done (e.g., constant versus variable temperature and the time scale of temperature variation, if any).

To explore the effect of temperature variation we modelled temperature using a sinusoidal function as follows:

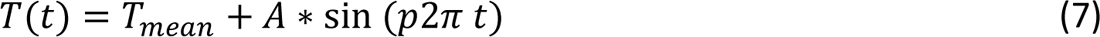

Where *T_mean_*is the average temperature, *A* represents the amplitude of temperature variation, and *p* is the period of forcing.

### Equilibrium behaviour along a temperature gradient

We begin by deriving the thermal performance curve (TPC) for algae (B), given by the per-capita growth rate (dB/Bdt) when nutrients are non-limiting. In this case, *N*/(*N* + *N*_0_) → 1 and the maximum equilibrium nutrient quota, Q, value, hereafter referred to as Q_max_, is reached. This maximum equilibrium value is obtained by simplifying Equation (3) and solving:

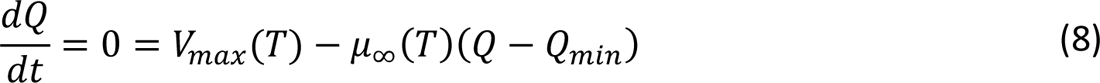

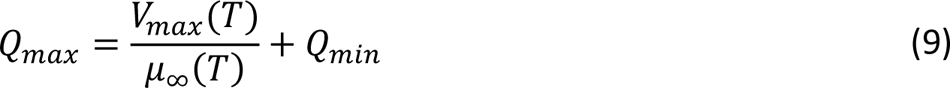

Substituting this value into Equation 5 yields the population’s fundamental TPC:

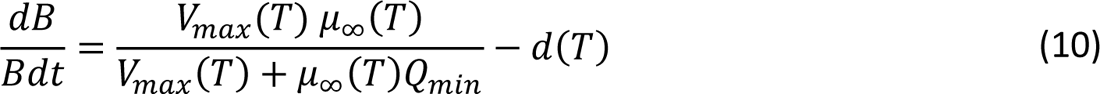

This curve represents a typical left-skewed unimodal function of temperature, which peaks at the temperature that maximizes growth (*T_opt_*), and positive growth rates are bounded by the lower and upper thermal limits, *T_min_*and *T_max_*, that define the fundamental thermal niche (Figure 1 and 2). The shape of this curve results from the fact that the first term of Equation 10 is unimodal with respect to *T*, reflecting the product of the two gaussian functions *V_max_*(*T*) and *μ_∞_*(*T*) (and therefore a symmetric, nearly gaussian function when *T_opt,μ_ = T_opt,v_*). Subsequently, subtracting *d*(*T*) from this curve creates the classic-shaped TPC (see (Amarasekare & Savage 2012; Vinton & Vasseur 2022), and results in *T_opt_ < T_opt,v_ & T_opt,μ_* because of this differential.

Previous work has established that the optimum for thermal performance (here measured by the per-capita growth rate dB/Bdt, decreases under nutrient limitation due to the non-linearity of the two terms in Equation 10 (Thomas *et al*. 2017; Vinton & Vasseur 2022). Under limiting nutrients, N cannot be factored out of Equation 10 (though it retains the same general shape) and nutrient limitation therefore scales this curve. We demonstrate this result in Figure 2B by solving dB/Bdt (from Equation 5) for different levels of nutrient uptake half-saturation constant (*N_0_*; indicating the efficiency of nutrient uptake, or de-facto nutrient limitation) where N is held at the supply concentration (*N_in_*=1) (i.e., the realized TPC). Eventually, nutrients become so limiting that the upper and lower limiting values of the thermal niche converge upon a single temperature; further nutrient limitation beyond this does not support population growth at any temperature.

With the addition of a dynamic nutrient pool (N, Equation 1), we can solve for the equilibrium algal biomass at different temperatures. This equilibrium is similar to the concept of the carrying capacity in the logistic growth model, only here it is an emergent property of the dynamics of our model (whereas in the logistic equation it is a parameter or input function; see (Vasseur 2020). We find that temperature and nutrients have an interactive effect on equilibrium biomass that does not match the thermal performance curve (Figure 2). Although equilibrium biomass is a unimodal function of temperature with lower and upper critical points matching those of the TPC, it is maximized at temperatures less than *T_opt_.* That is, the temperature response of equilibrium biomass, hereafter referred to as *K*(*T*) or the **thermal biomass curve** (TBC), is skewed opposite to that of the TPC. The difference between these two temperature optima, which we will denote *T_r_* and *T_K_*for optimal temperatures for growth and biomass, respectively, is affected by the accessibility of nutrients. Increasing the efficiency of nutrient uptake (i.e., decreasing *N_0_*) effectively decreases nutrient limitation and enhances the differences between the shape and optima of the thermal performance and thermal biomass curves (Figure 2). Here, we see that increasing uptake efficiency alters both curves such that they become more dissimilar when nutrients are non-limiting (i.e., saturated). On the other hand, when nutrients are highly limiting, the thermal performance and biomass curves ultimately converge to the same at the very point where *T_min_* and *T_max_* intersect; *T_r_*and *T_K_* necessarily converge. That is, when nutrients become so limiting that the TPC has only a single critical point (where dB/Bdt=0) the two curves have the same optimal temperature.

Importantly, this result suggests that population productivity (i.e., growth rate, r) and biomass (i.e., carrying capacity, *K*) have different temperature-dependencies, and that these differential responses are mediated (specifically, decreased) by nutrient limitation (Figure 2). Importantly, equilibrium biomass is always optimized at seemingly sub-optimal temperatures, relative to the TPC. We will hereafter refer to this as ***r-K* mismatch**, or simply mismatch for our purposes.

Although our modified Droop model is specific to phytoplankton growth, it is interesting to note that the mismatch between the thermal performance and thermal biomass curves has been shown to occur in similar models where nutrient quotas were not included as a dynamic component (Vinton and Vasseur 2022), suggesting that it is a general phenomenon generated by the interaction between temperature and nutrient (or, more broadly, resource) consumption and supply. In **Box 1**, we demonstrate the conditions under which our model can be simplified into a more general 2-equation system (analogous to Vinton and Vasseur’s consumer-resource model) and show that biomass is always maximized at lower temperatures than productivity (i.e., *T_K_* < *T_r_*) in this simpler model because of trade-offs between resource availability, (over-) consumption (i.e., density-dependence), and temperature-dependent death rates, which together regulate the “efficiency” of the system for turning resources into biomass. Specifically, at colder temperatures, a single unit of resources can support more population biomass because less is lost to death (or respiration) since *d*(*T*) is small. However at warmer temperatures, a single unit of resources supports less population biomass because *d*(*T*) is large. This observation combines with the fact that resource (nutrient) equilibrium densities follow a flat-bottomed U-shaped function of temperature; that is, resource equilibrium changes very little over much of the thermal niche, but the amount of consumers supported by it changes quite drastically (Figure B1). Therefore, the relationships identified in Box 1 ought to be generally true in consumer-resource models that fit our two simplifying assumptions (e.g., the model used in Vinton and Vasseur 2022).

### Box 1

#### What drives the thermal mismatch between production and biomass?

Figure 2 shows that equilibrium biomass consistently peaks at a lower temperature than thermal performance (per-capita population growth). The difference between the two peaks grows when individuals are more efficient at accessing their resources (nutrients in our case; decreasing *N0* of the uptake function). Vinton and Vasseur (2022) showed the same pattern when investigating the interactive nature of temperature- and resource-dependence in heterotrophic consumer populations, suggesting our inferences here are relevant for general consumer-resource systems. Here we investigate the origin of this mismatch. We find that the population’s equilibrium response to temperature is dependent on trade-offs between nutrient uptake/assimilation and turnover – both temperature-dependent processes – such that biomass optimization becomes less dependent on optimal growth conditions as nutrients become saturated.

In both models (ours and Vinton and Vasseur’s), equilibrium biomass is an asymmetric unimodal function of temperature that is skewed opposite to that of the TPC, such that equilibrium densities peak at values closer to *T_min_* than *T_max_* (i.e., the optimal temperature for biomass, *T_K_*, is less than the optimal temperature for the TPC, hereafter referred to as *T_r_*) (Figures 2, B1).

**Figure B1:**
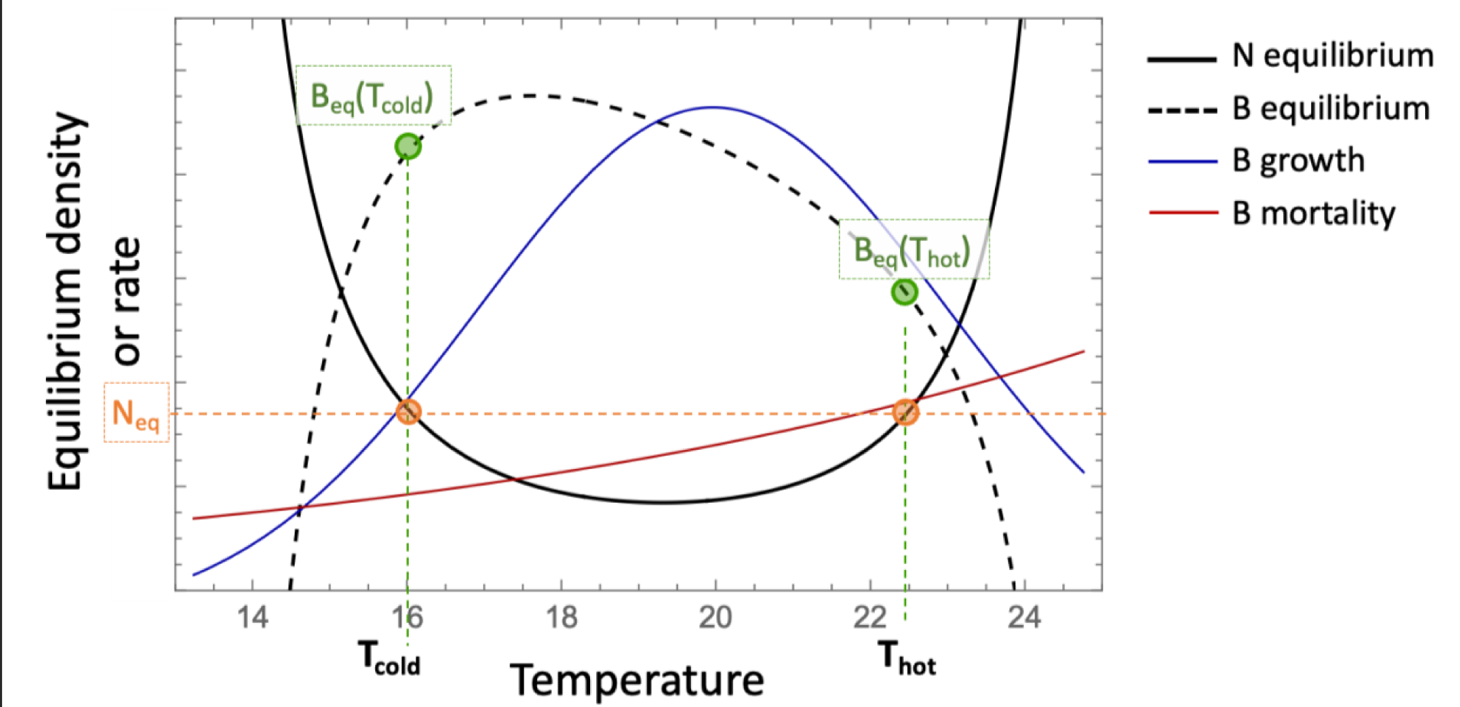
Summary of differential temperature responses of model equilibrium and vital rates.

The analytical derivation of the (consumer) population’s equilibrium biomass depends directly on the temperature dependent rate *V_max_*(*T*) (uptake or consumption), and on the equilibrium nutrient (resource) density which also changes as a function of temperature. From Equation 1, this expression is given by:

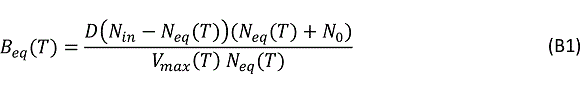

Where N_eq_(*T*) is the temperature dependent nutrient equilibrium. Direct inference from this expression is challenging due to the incorporation of both direct and indirect impacts of temperature. Instead, we employ two simplifications to understand the r-K mismatch.

First, given that the simpler 2-equation model of Vinton and Vasseur admits the same behavior as our more complex 3-equation model, we can recover the simpler 2-equation model of Vinton and Vasseur by assuming that the nutrient quota remains at equilibrium (Q = Q_eq_) and that Q_eq_ is constant across temperature. This effectively removes the dynamic effects of the nutrient Quota from our model and treats it as a constant. Therefore, from Equation 3:

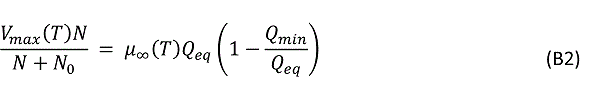

Substituting this relationship into Equation 5 gives:

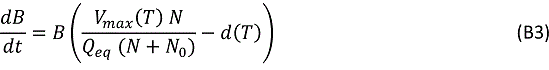

where 1/Q_eq_, when treated as a constant, is equivalent to the energy conversion efficiency in classical consumer-resource models. This assumption makes our approach analogous to classical consumer-resource models without a dynamic nutrient quota. In reality, different environmental or physiological conditions make this assumption more or less accurate – such as, for example, the availability of or efficiency of consuming nutrients (Figure 2).

Solving Equation B3 = 0, with some rearranging, yields:

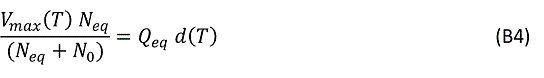

which can be substituted into expression B1 to give a simplified expression for the biomass equilibrium:

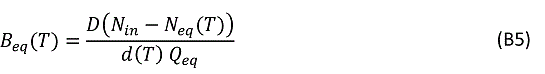

Notably, this is a simpler form than expression B1, but still retains both direct and indirect impacts of temperature change.

Our second simplifying assumption relies on the fact that the nutrient equilibrium is a nearly symmetric U-shaped function of temperature across the domain of the population’s fundamental niche. This means that we can identify two temperatures within this domain (*T_cold_* and *T_hot_*) that have equal nutrient equilibria (N_eq_(*T_cold_*) = N_eq_(*T_hot_*)), but different population biomass equilibria (B_eq_(*T_cold_*) ≠ B_eq_(*T_hot_*)) (Figure B1). At these two temperatures we can therefore remove the indirect effect of temperature via N_eq_ from equations B1 and B5.

Therefore, from these symbolic solutions (B1 and B5), differences between B_eq,hot_ and B_eq,cold_ are inversely proportional to the differences in uptake (*V_max_*(*T*)) and mortality (*d*(*T*)) rates at the two temperatures. That is, where N_eq,cold_ = N_eq,hot_ and B_eq,cold_ > B_eq,hot_, *V_max_*(*T_cold_*) < *V_max_*(*T_hot_*) and *d*(*T_cold_*) < *d*(*T_hot_*) (Figure B1). Specifically,

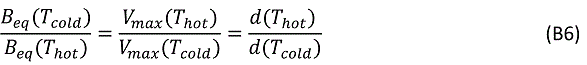

Despite the rate of nutrient uptake (*V_max_*) being lower at *T_cold_*, this is counteracted by the similarly lower mortality (i.e., turnover) rates and allows for larger biomass accrual. At *T_hot_*, nutrients are taken up at a faster rate, but more biomass is lost to turnover. Stronger temperature-responses of *V_max_*(*T*) and *d*(*T*) will cause larger differences between B_eq,hot_ and B_eq,cold_, and therefore stronger asymmetry in the equilibrium solution and increased mismatch between the optimal temperatures for biomass (equilibrium) and productivity (growth). Since *N_0_* scales the resource uptake rate, *V_max_* is maximized and it becomes intuitive why resource saturation (decreased *N_0_*) increases mismatch.

To summarize, for a given nutrient density (N_eq_), more population biomass can accumulate at lower temperatures, even though the nutrient uptake rate is lower. In other words, for a population that can achieve a particular N_eq_ (R* *à la* (Tilman 1982) for resources in general) but has a large *V_max_*, it will necessarily have fewer individuals because each of those individuals has a large impact on the nutrient levels. That is, the per capita impact of each B is higher for a higher *V_max_*, so there can be fewer consumptive individuals for a given N*_eq_. If a population can achieve the same N_eq_ but with a smaller *V_max_*, then each individual B will have a smaller effect on nutrients, allowing for more of them. Ultimately, the nutrient equilibrium is set by the ratio of *V_max_*(*T*)/*d*(*T*), but the biomass equilibrium is set by the magnitude of *V_max_*(*T*) and *d*(*T*).

Our full 3-dimensional model that incorporates a dynamic, temperature-dependent nutrient Quota adds complexity, yet retains the same general rules as identified in Box 1. From equation B5 (Box 1) we can see that the Quota is indeed important in determining equilibrium biomass, and therefore scales the ratios identified in solution B6 with temperature, providing important mechanistic nuance for understanding growth and biomass accumulation specific to phytoplankton populations. From Figures 1 and 2, temperature and nutrient limitation have an interactive effect on population biomass that is mediated by trade-offs between uptake and mortality (Box 1) and regulated via the nutrient Quota. When nutrients are saturated, growth occurs as soon as temperatures allow. This means that population biomass can be optimized at lower temperatures – where the equilibrium quota is larger (see Figure 2B for equilibria), and nutrients are efficiently turned into biomass without the large loss of biomass due to turnover at higher temperatures. Here, despite the surplus of resources fueling ample biomass, the rates of uptake and assimilation at these temperatures are lower, creating a greater lag through the Quota and overall slower growth and dynamics. Alternatively, when nutrients are limited *T_r_* and *T_K_* begin to converge. The fundamental niche shrinks (*T_max_* - *T_min_*) and low-temperature rates of uptake and assimilation are not sufficient to maximize population growth, so biomass is optimized at temperatures closer to where growth is optimized (i.e., *T_r_*) – reducing mismatch. However, biomass can not reach the same maximum as under nutrient-saturated conditions because of the corresponding increased turnover at these temperatures. Here, biomass is maximized under high flux (i.e., fast) conditions, with nutrients rapidly converted into population biomass – with less lag through the Quota (and less stored nutrients) – and then much of it lost to mortality.

It is worth noting that here we are assuming uptake and assimilation rates have symmetrical and equal thermal responses. Given the importance of trade-offs between uptake and assimilation rates in regulating the flux through the Quota, it is possible that asymmetry in these rates’ temperature responses may alter our results. However, while asymmetry between these temperature responses affects the magnitude of the biomass response, it does not affect mismatch because it simultaneously shifts the entire thermal niche to follow the temperature-dependence of nutrient uptake (see Appendix Figures A4-5). This is because uptake and assimilation are sequential processes, so do not have equal weighting in terms of regulating the thermal dependence of population growth and dynamics. That said, asymmetry in these temperature-responses (particularly when the thermal optimum for uptake is less than that for assimilation) allows for greater and earlier quota peaks, resulting in more efficient conversion to biomass before the quota is over-depleted by less efficient consumption at higher temperatures (Figure A5).

Importantly, the Quota represents the *potential* for biomass growth of a population, and therefore ought to be an important component regulating non-equilibrium responses to variable environments. The differential rates of biomass growth, and ability for nutrients to accumulate within the cell, across the thermal niche will become important under varying environments, such that both the nutrient storage ability (when assimilation rates are low) and implicit lag caused by the Quota ought to act as a buffering mechanism for populations during stressful or harsh times. Collectively, the nutrient quota – or more accurately the lag associated with the quota – ought to drive differential dynamic responses to environmental variability at high and low temperatures.

### *r-K* mismatch and implications for transient dynamics

We can now build off this insight on the equilibrium response to temperature, to understand transient population dynamics. Here, the temperature-dependence of vital rates and equilibrium biomass play important roles in determining the eventual state of a population (i.e., the equilibrium) and how long it takes to get there (i.e., transient length or return time). For example, while temperatures that optimize rates of population growth will result in short transients and therefore the fastest approach to equilibrium, this equilibrium is not optimized. Alternatively (and as suggested in Figure 2), temperatures that maximize equilibrium biomass (i.e., *K*) will lead to the highest population levels but take much longer to get there due to the lower rates at these temperatures. As described above, and eloquently shown by Anderson et al. (unpublished), despite the nutrient quota not being a strong determinant of equilibrium biomass, it can have a strong impact on the non-equilibrium dynamics of the system.

Furthermore, the fact that temperature cannot optimize both performance (growth) and biomass (production) has implications for population trajectories under climate change and for effective management strategies.

This suggests the intriguing possibility that certain forms of temperature variation between these two optima may be able to facilitate both optimal growth and optimal biomass. In classical population models (e.g., the logistic), *r* and *K* have an interesting relationship during dynamic population growth where the impact of *r* on dynamics is large when biomass is low, but weak when near *K*. This suggests that temperature could change dynamically to favour fast growth when biomass is low, and then favour large *K* when biomass has increased. Our model elucidates growth and biomass dynamics for phytoplankton populations more mechanistically than the logistic model, but follows the same principles. *T_r_* maximizes population growth at near-0 biomass densities, but as biomass grows and resources become more depleted, this temperature may no longer be the “optimal” environment for a population collectively to be in. At higher densities, it becomes more beneficial for temperatures to be lower (near *T_K_*) where the efficiency of conversion from nutrients to biomass is maximized, turnover is low, and therefore biomass can be maximized. Indeed, we can see that *T_r_*and *T_K_* result in both different equilibria and transient lengths, but when properly timed, a transition between the two temperatures can maximize both the rate of population increase and ultimate biomass obtained (Figure 3). These differential responses of population growth and size at different temperatures also imply potentially interesting dynamic effects of continuously varying temperatures (e.g., seasonality).

**Figure 3.**
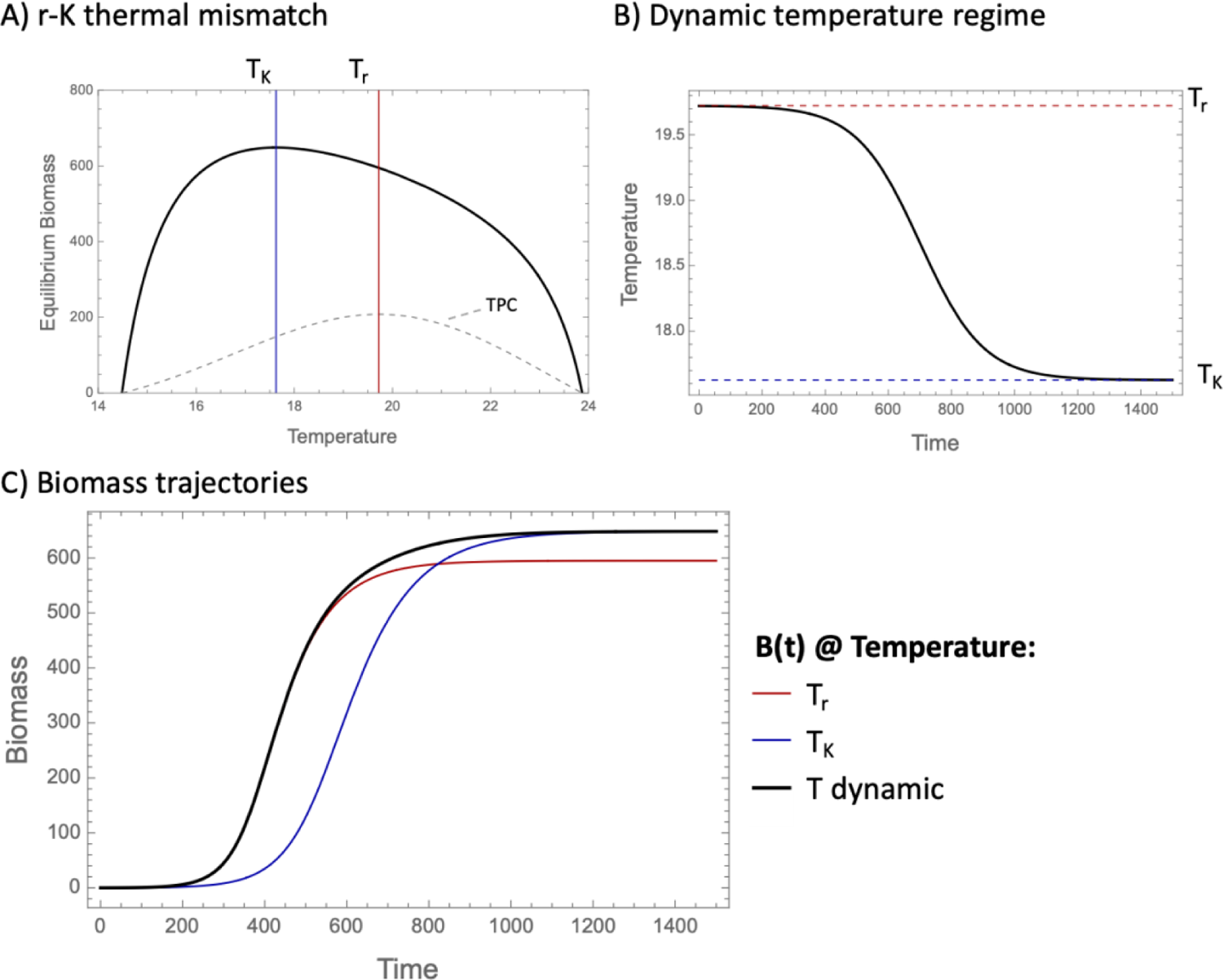
Thermal optima for maximizing biomass (K) or growth rate (r) and corresponding transient dynamics (approach to equilibrium). When temperature is varied such that temperature starts at T_r_ (maximum growth rates when biomass densities are low), then gradually decreases to T_K_ (maximum equilibrium biomass), the population is able to reach equilibrium faster. In this sense, temperature is optimized to facilitate rapid growth from low densities and keep this pace as it then approaches maximum biomass.

### Non-equilibrium dynamics and population responses

Figures 1-3 collectively show that population growth rate and quota-induced lags, transient lengths, and equilibrium biomass have different and interacting temperature-dependencies. This means that the time scales that population dynamics operate on are temperature dependent and suggests that the time scale of environmental variation ought to be important in determining long-term population dynamics. Effective population forecasting requires us to identify time scales at which variation in temperature is going have to important effects beyond those predicted by equilibrium dynamics or average temperatures.

When temperature is varied sinusoidally between the boundaries of a population’s thermal niche (*T_min_* and *T_max_*), the equilibrium approaches (but does not pass) 0 at both of these extreme temperatures – therefore doubling the period of expected biomass dynamics over one period of temperature forcing. Figure 4 shows the biomass dynamics over time under this form of temperature variation, relative to the (changing) equilibrium (also see Figure A7 in the Appendix for additional temporal scales and Figures A8-9 for dynamics of N and Q). Note that when the forcing period is *very* long (e.g., with this parameterization, >500 000 time steps), biomass almost perfectly tracks the equilibrium curve (Figure A7) and population dynamics can therefore be accurately predicted using the thermal biomass curve at all temperatures. At the other extreme, when forcing is very fast, biomass dynamics effectively cannot respond to the rapidly changing temperature and biomass becomes nearly invariant – approaching the mean equilibrium biomass over the thermal range (Figure 4C and Figure A7). These responses reflect two extremes of dynamical responses along a gradient of environmental forcing speed, relative to the underlying vital rates of the model (i.e., the population’s rate of change, or ability to respond to environmental changes). At intermediate speeds of temperature variation, the dynamics become less predictable (Figure 4). Here, there is a dynamic interplay between a changing attractor (equilibrium), local stability, and thermally asymmetric population rates of change (e.g., growth rates and the Quota-induced lag), together causing the dynamics to lag unevenly behind the changing deterministic equilibrium; we thus see the amplitude of biomass variation decreasing with increasing forcing speeds (Figure 4).

**Figure 4.**
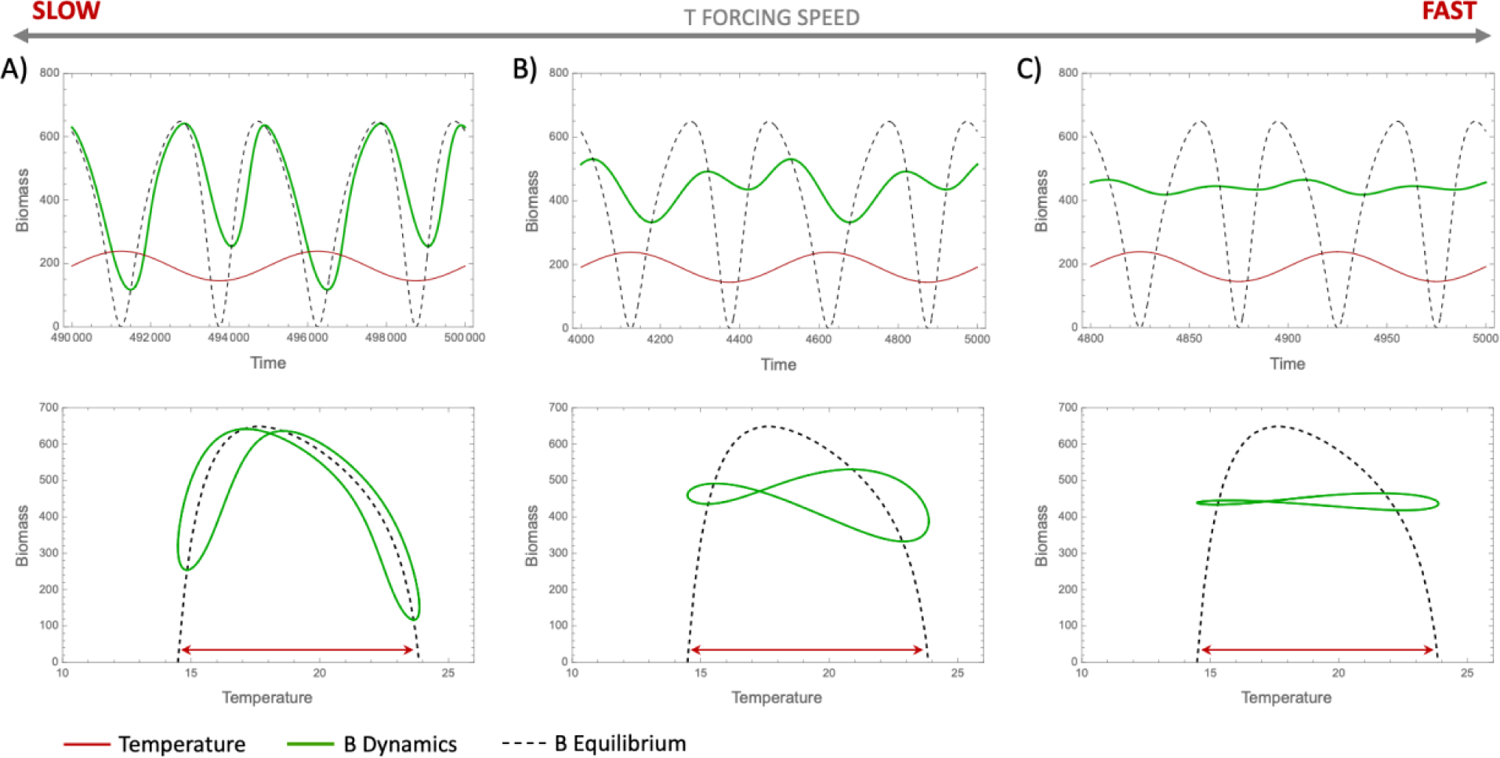
Population dynamics, relative to the temperature-dependent equilibrium, in response to sinusoidally varying temperatures between T_min_ and T_max_. Here, dynamics have differential abilities to “track” the changing equilibrium depending on the speed of forcing and temperature-dependent rates of population growth. Forcing periods shown here: A) 5000, B) 500, and C) 100 time units. Temperature in top row is scaled for visualization purposes (multiplied by 10).

Under still relatively slow forcing (e.g., Figure 4A), population dynamics can nearly track the equilibrium but fail near the temperature extremes – that is, where population rates of change slow and local stability approaches zero (Figure A6). This is also where thermal asymmetry in the Quota-induced lag becomes apparent: the lag is higher at low versus high temperatures, even where population growth rates (dB/Bdt) are equally low (i.e., near *T_min_* and *T_max_*), meaning the time scale that population dynamics operate on is different at high and low temperatures, and biomass dynamics are less responsive to changing temperature when they are low. What this means is that populations track the equilibrium better at high temperatures than at low temperatures, and are therefore more likely to collapse when temperatures approach *T_max_* (most apparent in Figure 4A) or even surpass the temperature extremes (e.g., see Figure A10). On the other hand, the more pronounced lag at lower temperatures means that the population will retain higher biomass for longer as temperatures approach *T_min_* (Figure 4, as well as Figure A10 for temperature varying beyond the thermal niche). This suggests that as long as forcing is fast enough, the population effectively does not have enough time to crash at these low temperatures before temperature rises again (Figure 4A,B). Next, as temperature forcing becomes faster (Figure 4B,C), this same dynamic thermal asymmetry remains, and the temperature-dependent rates of change drive a form of perpetual “overshoot” at high temperatures, reinforcing this high-temperature-variability (relative to low temperatures) even though the dynamics can no longer closely follow the equilibrium. The magnitude of this variability is again dependent on the time scale of temperature variation, relative to the population, eventually approaching invariant population dynamics as forcing speeds further increase.

Finally, the mechanisms behind these dynamics are qualitatively general despite a modest interactive effect with nutrient limitation (saturation of nutrient uptake, or “efficiency”) on the realized dynamics under variable temperature. When nutrient uptake is less efficient (i.e., *N_0_*is higher), both the TPC and the equilibrium response are altered (Figure 2), and together these changes reduce the asymmetric dynamics between high and low temperatures (i.e., via the quota) while also lowering the overall “pace” of population growth. Under these circumstances, *r-K* mismatch is decreased, the buffering effect of low temperatures seen above is lessened, and the population dynamics become more invariant than when nutrients are less limited (Figure A11). Note, however, that this is in large part a result of *relative* time scales of population dynamics and temporal forcing, since a given frequency of forcing will “seem” faster to a population with slower growth rates.

## Discussion

Here we demonstrate that under the assumption of a constant supply of resources (a chemostat supplying nutrients), the thermal performance of a phytoplankton population (i.e., the temperature dependence of per-capita growth rate under density-independent conditions) is different than the thermal performance of equilibrium biomass, with biomass always peaking at cooler temperatures than performance. The thermal biomass curve outlined here describes how carrying capacity (*K*) varies with temperature for a population consuming finite resources (with a constant supply); hence, we refer to this thermal differential as *r-K* mismatch.

Furthermore, we demonstrate that these thermal relationships have differential responses to changes in resource availability; the optimal temperatures for the two attributes converge as resources become more limiting, ultimately intersecting at the point where resources are so scarce that they can no longer support a viable population (Figure 2). Our mechanistic insight into the drivers of *r-K* mismatch for phytoplankton populations show that this pattern is a result of trade-offs between nutrient uptake and death, and that inclusion of a dynamic nutrient quota allows us to determine how both equilibrium and non-equilibrium dynamics depend on a combination of nutrients, temperature, and variability (speed of fluctuations) in the environment. Furthermore, these results – and the trade-offs generating them – ought to be common phenomenon for more general consumer-resource interactions (i.e., thermal mismatch between *r* and *K* in a population with density dependent growth; Box 1). Specifically, these patterns ought to always hold so long as mortality increases with temperature and differences between the thermal responses of growth and mortality define an organism’s thermal niche – together driving *r-K* mismatch. Indeed, although our model and analytical results here rely on a unimodal function of temperature for population growth (nutrient uptake and assimilation rates), we note that the model behaves similarly when we shift to monotonically increasing (i.e., exponential) functions (see Appendix, Figure A3). This is in line with common “double exponential” approaches to modelling population performance (e.g., (Thomas *et al*. 2017)), and importantly highlights that our results reveal a general representation of the thermal response of growth, turnover and density-dependence.

One of the major challenges associated with predicting the dynamic response of populations to changes in temperature and other environmental attributes is the role of indirect effects; temperature-dependent changes in the productivity of resources or density of competitors, for example, make it difficult to anticipate how a focal species will respond. At the population level, a variety of indirect effects manifest as changes in the strength of density dependence.

Understanding these changes will inform improved general models of population responses. Our demonstration of differences in the optimal temperatures for performance and biomass is generated by a strengthening of density-dependence as temperature increases, such that at warmer temperatures fewer individuals are supported per unit of resource (Box 1). When framed in terms of classic Logistic population dynamics, the increase in the strength of density-dependence leads to an *r-K* mismatch, where *r* is optimized at a warmer temperature than that which optimizes *K*. Importantly, the thermal dependence of both rates of change (productivity) and equilibrium (biomass) will dictate population trajectories and dynamics under global change, and understanding the potential for thermal mismatches is necessary to predict or forecast dynamics into the future.

Predictive population forecasts are often based on organisms’ physiological thermal performance (i.e., their TPCs extrapolated to match changing environments), but our results suggest that the effect of global change on population densities may not match these forecasts: when there is an *r-K* mismatch, projections cannot be informed by the TPC of fitness (*r*(*T*)) alone. Clearly TPCs are necessary for understanding rates of productivity for populations, energy flux within food webs and nutrient cycling within ecosystems, but without a mechanistic understanding of biomass responses we cannot accurately predict extinction risk of populations nor numerical responses of important processes (e.g., interactions, energy flux). Certainly, increasing temperatures towards *T_max_* – when *r* and *K* are both declining – will have strong effects on the extinction risk of a population, relative to any estimates that ignore any effect on *K*. However, we also show that there is a range of temperatures over which the TPC increases, but *K* decreases, where the risk of warming could be much harder to evaluate – for example due to increasing resilience (local stability or rates of “attraction”) toward a smaller equilibrium population size. Furthermore, the nonlinear effects of nutrient uptake saturation (i.e., effective nutrient limitation) on both the magnitude and shape of the thermal biomass curve highlights another layer of complexity important for accurate population forecasting under global change, and similarly suggests interesting implications for temporal patterns in population dynamics under variable temperature and nutrient regimes.

Elucidating the thermal biomass curve for populations is a necessary first step towards understanding population dynamics – and therefore extinction risk – under global change. These results provide a baseline for understanding the patterns we see in non-equilibrium dynamics under variable environments, even when the dynamics do not perfectly “track” the equilibrium. As shown here, the temporal dynamics exhibited by a species in a thermally varying environment indeed get interesting when variation in *r* and *K* occur simultaneously. In ‘classic’ models where only r responds to the environment, the effect of the environment quickly wanes as a population approaches its carrying capacity. However, in the case where both *r* and *K* continually change in response to the environment, there is an interplay among the two parameters: *K* sets the target to which the population is attracted, and the strength of that attraction is determined by *r*. In this case, it is important to understand where natural thermal variation lies relative to both the thermal biomass and performance curves. For example, in scenarios where thermal variability exists only below *T_r_* (the thermal optimum for growth) one might predict that populations will directly follow this environmental signature (e.g., (Smith 1997), despite the true period of population fluctuations being doubled if the variation actually spans either side of *T_K_* (that is, the actual attractor), leading to fundamentally different predicted population dynamics over time. As an example of this, phytoplankton are known to have spring and fall peaks in biomass, a phenomenon generally thought to be driven by a combination of nutrient cycling and predation (Cebrian & Valiela 1999; Martinez *et al*. 2011; Sigler *et al*. 2014), but which could in fact be enhanced by temperature in a case such as this.

Under varying thermal conditions, our results also highlight important thermal asymmetries in population rates of change and how populations respond to changing environments – a result that can be explained mechanistically by the implicit lag associated with our dynamic nutrient quota. While others have incorporated explicit lags into nutrient quota dynamics (e.g., (Cunningham & Nisbet 1980), thermal asymmetries within our purely monotonic model clearly have implications for population dynamics. In Figure 4, we showed that the time scale of environmental variation is important for determining population dynamics as the relative influence of the thermally asymmetric lag wanes with increasing forcing period. This in turn changes the potential for collapse when temperatures approach an organism’s thermal limits. Notably, these results suggest that 1) lagged population dynamics mean that populations can likely withstand brief periods with temperatures outside the fundamental niche; and 2) brief periods with temperatures below *T_min_* ought to be substantially less catastrophic for population persistence than brief periods above *T_max_* (Figures 4, A7 and A10). Furthermore, a population’s response to or recovery from perturbations (i.e., mass mortality events or environmental stochasticity) based on its rates of change and lagged responses has important implications for its dynamics in variable environments and the potential to detect warning signs of collapse.

These results indicate that classical approaches to detecting early warning signals of critical transitions (e.g., critical slowing down) may be impeded by the interacting thermal asymmetries of growth rates, equilibria, and lagged dynamics. Finally, the non-equilibrium dynamics here highlight an important relationship between temperature and population variability resulting from these thermal asymmetries (Figure 4). These results suggest that periodic environments (e.g., seasonal) could lead to increased variability in warmer (average) climates, and similarly that we may see more variability during warm (summer) versus cool (winter) times. This also suggests implications for population dynamics – and primary productivity for whole ecosystems – when seasons become less predictable (i.e., variability in the environment is amplified by higher vital rates).

Importantly, linking physiological processes at the individual level to higher order processes and dynamics at the population-level allows us to more intentionally build generalizable population models that are better grounded in first principles. There is often a need to simplify to more general models and contexts when making predictions and forecasting, and for developing fundamental ecological theory. One such example is the Logistic model, which continues to be central to the study of population dynamics, despite the reliance of parameters on environmental attributes (e.g., resource limitation and temperature) remaining open to interpretation. Investigating more specific models allows us to have a more reasonable understanding of the temperature-dependence of important rates and processes. Here, we have developed important mechanistic understanding of the dynamics of phytoplankton populations – the keystone to energy supply in all aquatic food webs, central to global carbon cycling, and a common study taxon for linking theoretical and empirical approaches in ecology. Simultaneously, our analytical insights apply to more generalizable model contexts with the goal of constructing fundamental theory in an intentional, informed way. Specifically, our r-K mismatch provides a framework for the nutrient- and temperature-dependence of population dynamics, and the next step ought to be developing a generalizable analytical form consistent with both our model and that used by Vinton and Vasseur (2022). Gaining a better understanding of the “true” shape of the thermal biomass curve, *K*(*T*), will be important for understanding the thermal response of primary production for whole food webs and therefore general ecological functioning in changing environments.

## Acknowledgements

This work was financially supported by the Yale Center for Natural Carbon Capture, and CB was funded through a NSERC PDF. We would like to thank David Anderson, Colin Kremer, Sam Fey, and Alison Robey for feedback on an earlier draft of this manuscript.

## References

Aksnes, D. & Egge, J. (1991). A theoretical model for nutrient uptake in phytoplankton. Mar. Ecol. Prog. Ser., 70, 65–72.

Amarasekare, P. (2015). Effects of temperature on consumer-resource interactions. J Anim Ecol, 84, 665–679.

Amarasekare, P. & Savage, V. (2012). A Framework for Elucidating the Temperature Dependence of Fitness. The American Naturalist, 179, 178–191.

Bernhardt, J.R., Sunday, J.M. & O’Connor, M.I. (2018a). Metabolic Theory and the Temperature-Size Rule Explain the Temperature Dependence of Population Carrying Capacity. The American Naturalist, 192, 687–697.

Bernhardt, J.R., Sunday, J.M., Thompson, P.L. & O’Connor, M.I. (2018b). Nonlinear averaging of thermal experience predicts population growth rates in a thermally variable environment. Proc. R. Soc. B., 285, 20181076.

Bestion, E., Schaum, C. & Yvon-Durocher, G. (2018). Nutrient limitation constrains thermal tolerance in freshwater phytoplankton. Limnol Oceanogr Lett, 3, 436–443.

Brown, J.H., Gillooly, J.F., Allen, A.P., Savage, V.M. & West, G.B. (2004). Toward a Metabolic Theory of Ecology. Ecology, 85, 1771–1789.

Cebrian, J. & Valiela, I. (1999). Seasonal patterns in phytoplankton biomass in coastal ecosystem. Journal of Plankton Research - J PLANKTON RES, 21, 429–444.

Cunningham, A. & Nisbet, R.M. (1980). Time Lag and Co-operativity in the Transient Growth Dynamics of Microaigae. Journal of Theoretical Biology, 84, 189–203.

Dillon, M.E., Woods, H.A., Wang, G., Fey, S.B., Vasseur, D.A., Telemeco, R.S., et al. (2016). Life in the Frequency Domain: the Biological Impacts of Changes in Climate Variability at Multiple Time Scales. Integr. Comp. Biol., 56, 14–30.

Doney, S.C., Ruckelshaus, M., Emmett Duffy, J., Barry, J.P., Chan, F., English, C.A., et al. (2012). Climate Change Impacts on Marine Ecosystems. Annual Review of Marine Science, 4, 11– 37.

Droop, M.R. (1974). The nutrient status of algal cells in continuous culture. J. Mar. Biol. Ass., 54, 825–855.

Droop, M.R. (1977). An Approach to Quantitative Nutrition of Phytoplankton. The Journal of Protozoology, 24, 528–532.

Englund, G., Öhlund, G., Hein, C.L. & Diehl, S. (2011). Temperature dependence of the functional response. Ecology Letters, 14, 914–921.

Fey, S.B., Kremer, C.T., Layden, T.J. & Vasseur, D.A. (2021). Resolving the consequences of gradual phenotypic plasticity for populations in variable environments. Ecological Monographs, 91, e01478.

Fussmann, K.E., Schwarzmüller, F., Brose, U., Jousset, A. & Rall, B.C. (2014). Ecological stability in response to warming. Nature Clim Change, 4, 206–210.

Gillooly, J.F., Brown, J.H., West, G.B., Savage, V.M. & Charnov, E.L. (2001). Effects of Size and Temperature on Metabolic Rate. Science, 293, 2248–2251.

Grover, J.P. (1992). Constant- and variable-yield models of population growth: Responses to environmental variability and implications for competition. Journal of Theoretical Biology, 158, 409–428.

Hastings, A., Abbott, K.C., Cuddington, K., Francis, T., Gellner, G., Lai, Y.-C., et al. (2018). Transient phenomena in ecology. Science, 361, eaat6412.

Higgins, K., Hastings, A., Sarvela, J.N. & Botsford, L.W. (1997). Stochastic Dynamics and Deterministic Skeletons: Population Behavior of Dungeness Crab. Science, 276, 1431– 1435.

Huey, R.B. & Kingsolver, J.G. (2019). Climate Warming, Resource Availability, and the Metabolic Meltdown of Ectotherms. The American Naturalist, 194, E140–E150.

Jarvis, L., McCann, K., Tunney, T., Gellner, G. & Fryxell, J.M. (2016). Early warning signals detect critical impacts of experimental warming. Ecology and Evolution, 6, 6097–6106.

Layden, T.J., Kremer, C.T., Brubaker, D.L., Kolk, M.A., Trout-Haney, J.V., Vasseur, D.A., et al. (2022). Thermal acclimation influences the growth and toxin production of freshwater cyanobacteria. Limnology and Oceanography Letters, 7, 34–42.

Lemoine, N.P. (2019). Considering the effects of temperature × nutrient interactions on the thermal response curve of carrying capacity. Ecology, 100.

León, J.A. & Tumpson, D.B. (1975). Competition between two species for two complementary or substitutable resources. Journal of Theoretical Biology, 50, 185–201.

Martinez, E., Antoine, D., D’Ortenzio, F. & de Boyer Montégut, C. (2011). Phytoplankton spring and fall blooms in the North Atlantic in the 1980s and 2000s. Journal of Geophysical Research: Oceans, 116.

McCoy, M.W. & Gillooly, J.F. (2008). Predicting natural mortality rates of plants and animals. Ecology Letters, 11, 710–716.

Nisbet, R.M. & Gurney, W.S.C. (1982). *Modelling fluctuating populations*. Wiley, Chichester; New York.

Sauterey, B. & Ward, B.A. (2022). Environmental control of marine phytoplankton stoichiometry in the North Atlantic Ocean. Proc. Natl. Acad. Sci. U.S.A., 119, e2114602118.

Savage, V.M., Gillooly, J.F., Brown, J.H., West, G.B. & Charnov, E.L. (2004). Effects of Body Size and Temperature on Population Growth. The American Naturalist, 163, 429–441.

Sigler, M.F., Stabeno, P.J., Eisner, L.B., Napp, J.M. & Mueter, F.J. (2014). Spring and fall phytoplankton blooms in a productive subarctic ecosystem, the eastern Bering Sea, during 1995–2011. Deep Sea Research Part II: Topical Studies in Oceanography, 109, 71– 83.

Slein, M.A., Bernhardt, J.R., O’Connor, M.I. & Fey, S.B. (2023). Effects of thermal fluctuations on biological processes: a meta-analysis of experiments manipulating thermal variability. Proceedings of the Royal Society B: Biological Sciences, 290, 20222225.

Smith, H.L. (1997). The periodically forced Droop model for phytoplankton growth in a chemostat. Journal of Mathematical Biology, 35, 545–556.

Smith, H.L. & Waltman, P. (1994). Competition for a Single Limiting Resource in Continuous Culture: The Variable-Yield Model. SIAM Journal on Applied Mathematics, 54, 1113– 1131.

Steffen, W., Broadgate, W., Deutsch, L., Gaffney, O. & Ludwig, C. (2015). The trajectory of the Anthropocene: The Great Acceleration. The Anthropocene Review, 2, 81–98.

Thomas, M.K., Aranguren-Gassis, M., Kremer, C.T., Gould, M.R., Anderson, K., Klausmeier, C.A., et al. (2017). Temperature–nutrient interactions exacerbate sensitivity to warming in phytoplankton. Glob Change Biol, 23, 3269–3280.

Tilman, D. (1982). Resource competition and community structure. Monographs in population biology. Princeton University Press, Princeton, N.J.

Uszko, W., Diehl, S., Englund, G. & Amarasekare, P. (2017). Effects of warming on predator–prey interactions – a resource-based approach and a theoretical synthesis. Ecology Letters, 20, 513–523.

Vasseur, D.A. (2020). The impact of temperature on population and community dynamics. Theoretical ecology: concepts and applications, 243–262.

Vasseur, D.A., DeLong, J.P., Gilbert, B., Greig, H.S., Harley, C.D.G., McCann, K.S., et al. (2014). Increased temperature variation poses a greater risk to species than climate warming. Proc. R. Soc. B., 281, 20132612.

Vinton, A.C. & Vasseur, D.A. (2022). Resource limitation determines realized thermal performance of consumers in trophodynamic models. Ecology Letters, 25, 2142–2155.

