## Appendix for "Interactions between temperature and nutrients determine the population dynamics of primary producers"

### Supplementary material for “Interactions between temperature and nutrients determine the population dynamics of primary producers”

Carling Bieg[1]* and David Vasseur[1]

1. Ecology and Evolutionary Biology, Yale University

#### Appendix

##### Equilibrium solutions & isoclines

The full equilibrium solutions including temperature effects are too complicated to be useful symbolically, however we can simplify the temperature-dependent components in the following way to show the general equilibrium solutions when population persistence is allowed. Thus, we leave the rates and state variables as temperature-dependent functions within the solutions:

$$B_{eq}(T) = \frac{D\big(N_{in} - N_{eq}(T)\big)\big(N_{eq}(T) + N_0\big)}{V_{max}(T)\,N_{eq}(T)} \tag{A1}$$

$$Q_{eq}(T) = \frac{Q_{min}\mu_{\infty}}{\mu_{\infty} - d(T)} \tag{A2}$$

$$N_{eq}(T) = -\frac{Q_{min}N_0\mu_{\infty}(T)d(T)}{d(T)V_{max}(T) + Q_{min}\mu_{\infty}(T)d(T) - \mu_{\infty}(T)V_{max}(T)} \tag{A3}$$

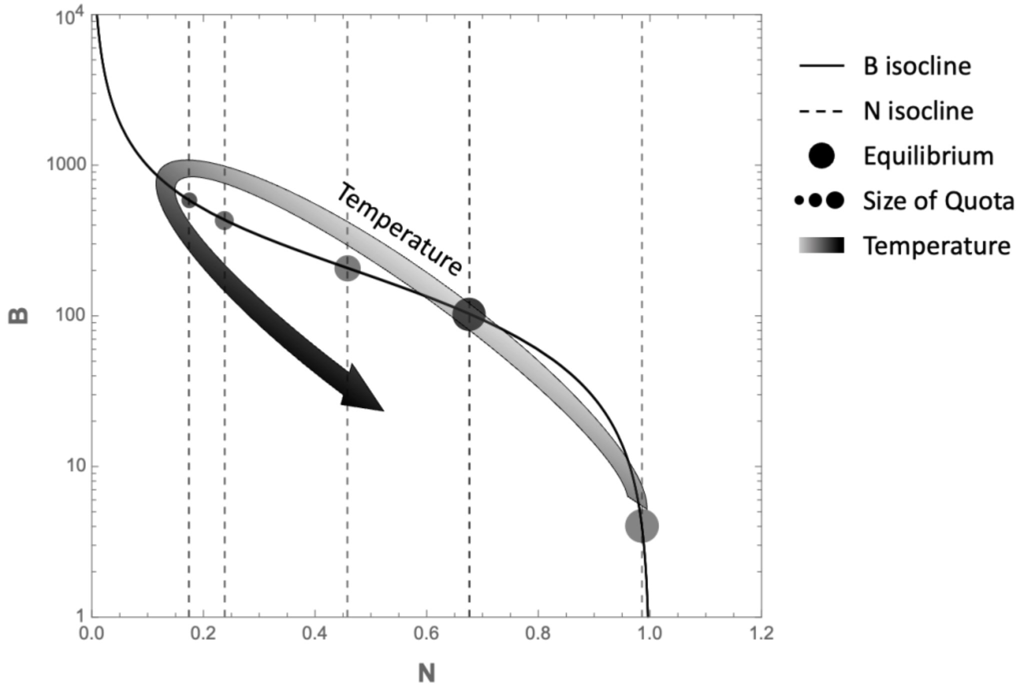


Figure A1. Effect of increasing temperature (opacity increases with temperature) in model isoclines and equilibrium. Equilibrium quota is reflected by the size of the circles marking the equilibrium. Temperatures shown are 14.5, 15.5, 17, 20 and 23.5, thus spanning the entire thermal niche.

**r-K mismatch generality: parameters and underlying assumptions of thermal responses**

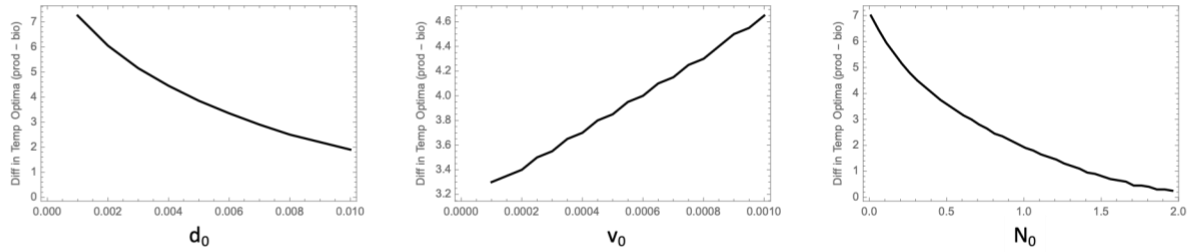


Figure A2: Effect of parameters on r-K mismatch. d0 affects turnover rate and v0 and N0 both affect accessibility of nutrients – all of which alter the potential for biomass accumulation and therefore the temperature response of biomass. When biomass cannot be accrued at sub-optimal temperatures, mismatch is decreased.

It is common for models of temperature-dependent phytoplankton growth to use a double-exponential approach, rather than the gaussian-exponential approach used here (e.g., (Thomas *et al.* 2017). That is, growth and loss (births and deaths) are both modelled as exponential functions of temperature, such that their intersection points determine the limits to an organism's fundamental niche. In both cases, the resulting TPC is a left-skewed unimodal function of temperature, however the shape of that curve may be slightly different depending on where the responses intersect. Here, we see that these assumptions do not fundamentally alter our results regarding the shape of the TBC (relative to the TPC) and thus the general result of r-K mismatch. We modelled nutrient uptake and assimilation as exponential functions of temperature and explored the effect on both thee TPC and TBC. While the analytical solution for the TBC slightly differs from ours, this is largely due to a differential scaling of temperature's effects, and we see that the results remain qualitatively unchanged.

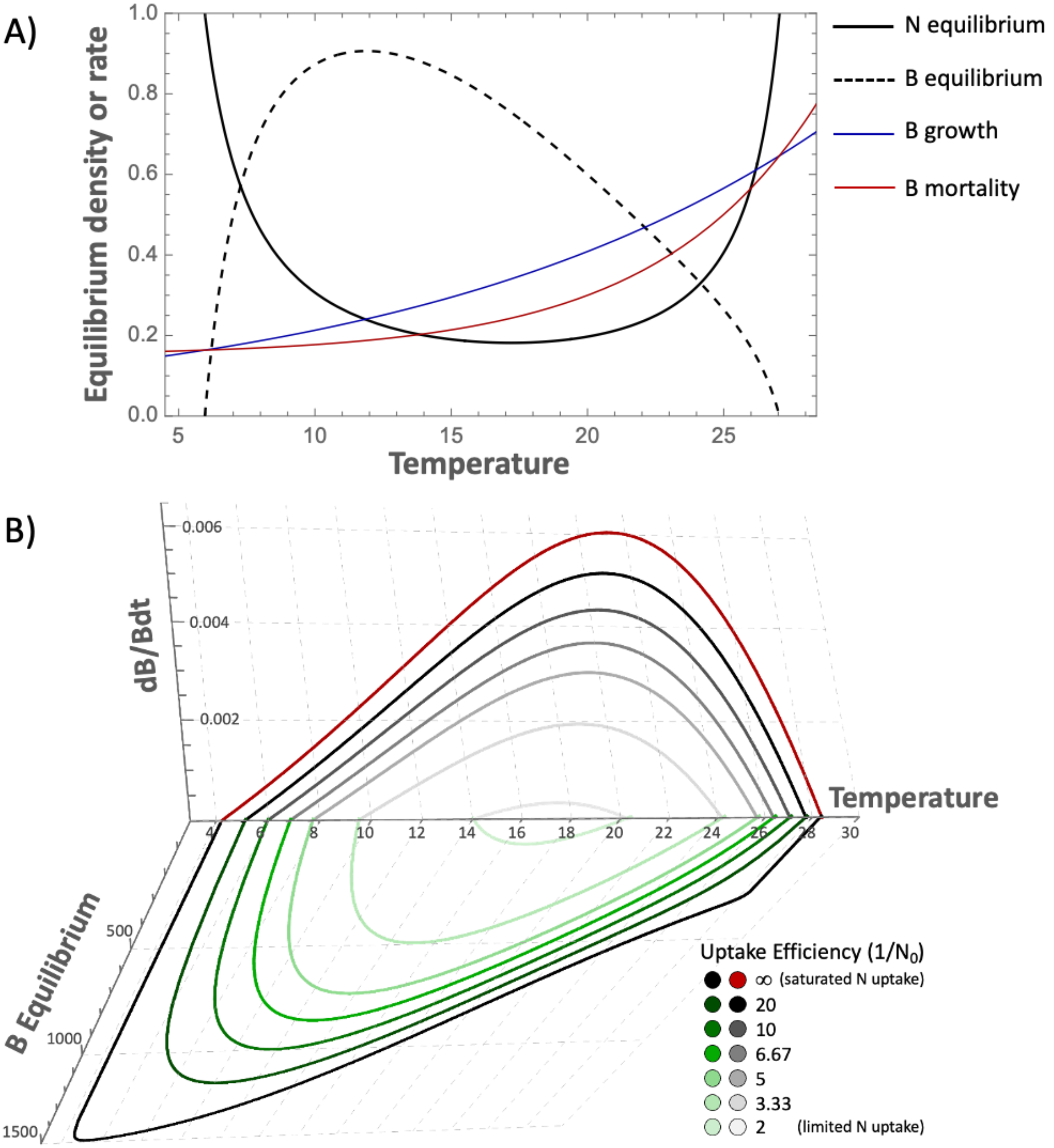


Figure A3. A) Temperature-dependent rates and thermal responses of biomass and growth, and B) the effect of nutrient limitation on these thermal responses.

**Asymmetry in temperature-dependent rates**

*Temperature-response asymmetry*

Asymmetry in optimal temperatures for uptake vs. assimilation (i.e., $T_{opt,v}/T_{opt,\mu}$ =/= 1) does not have a large effect on r-K mismatch.

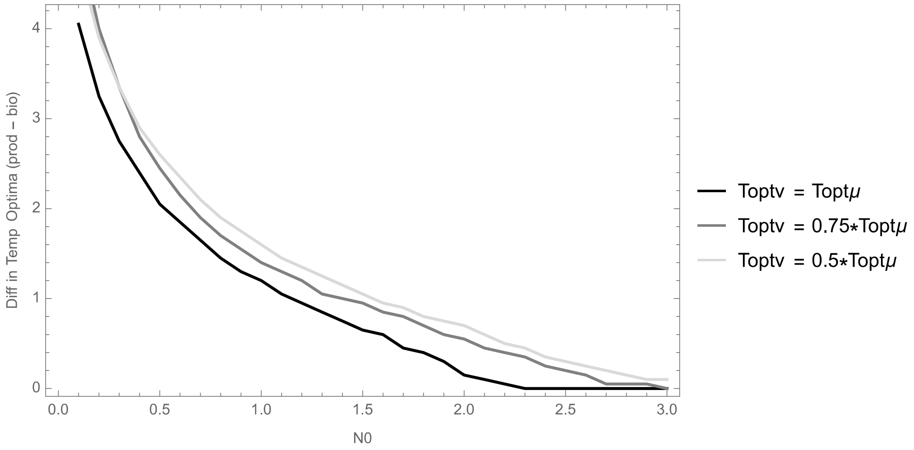


Figure A4: Asymmetry in temperature response of uptake vs. assimilation rates (i.e., differential optimal temperatures) has negligible effect on r-K mismatch across levels of $N_0$.

The biomass response curve is indeed affected by temperature-response asymmetry (Figure A5 a). Asymmetry (notably when uptake's optimal temperature, $T_{opt,v}$, is lower than assimilation's, $T_{opt,\mu}$) increases the amount of biomass accrual. However, mismatch is not affected by temperature-response asymmetry because this asymmetry also shifts – and broadens – the thermal niche, such that the TPC follows the temperature response of nutrient uptake ($T_{opt,v}$) (Figure A5 b). So mismatch is not affected by temperature-response asymmetry, but the amount of biomass accrual (i.e., max carrying capacity) is. In other words, if the optimal temperature for nutrient uptake is lower than that for assimilation, the whole thermal niche is shifted lower but the system as a whole becomes more efficient at turning resources into biomass.

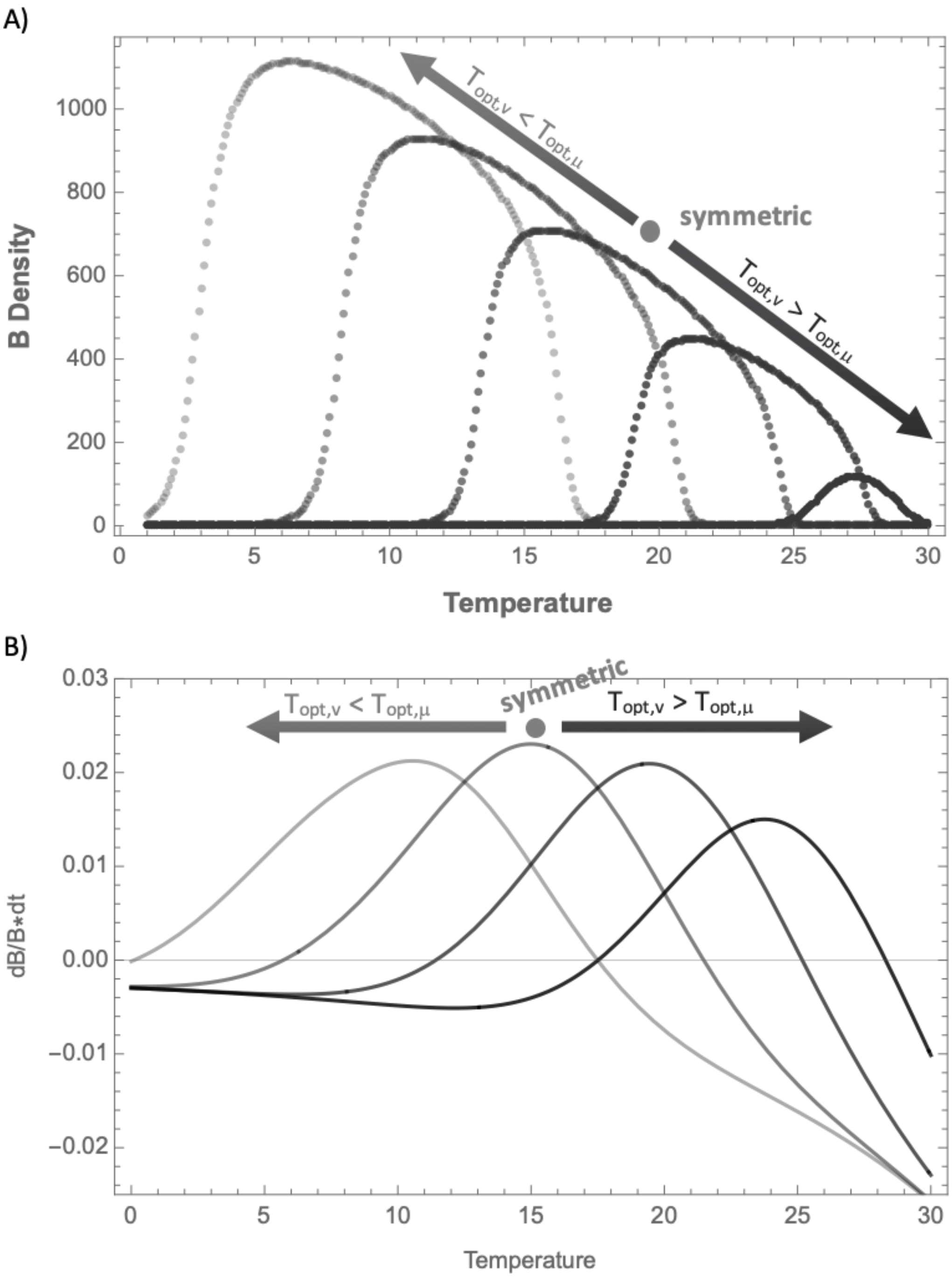


Figure A5. Effect of asymmetry in temperature responses of uptake (V) and assimilation ($\mu$) rates on A) equilibrium biomass, and B) dB/Bdt (i.e., the TPC).

**Variable environment & non-equilibrium dynamics**

*Local stability across thermal niche*

Note that these dynamics also make sense in light of the eigenvalue solutions, which are an artifact of population growth rate and therefore skewed in a similar fashion to the TPC (Figure A6). The non-trivial (interior) equilibrium retains near-maximal stability (most negative) at high temperatures such that the equilibrium will have a strong locally attracting force for a longer time (if we consider increasing temperature over time, approaching $T_{max}$). The more gradual decrease in stability with cooling temperatures means that the equilibrium will be less attractive as temperature cools and the system will be less likely to track this equilibrium.

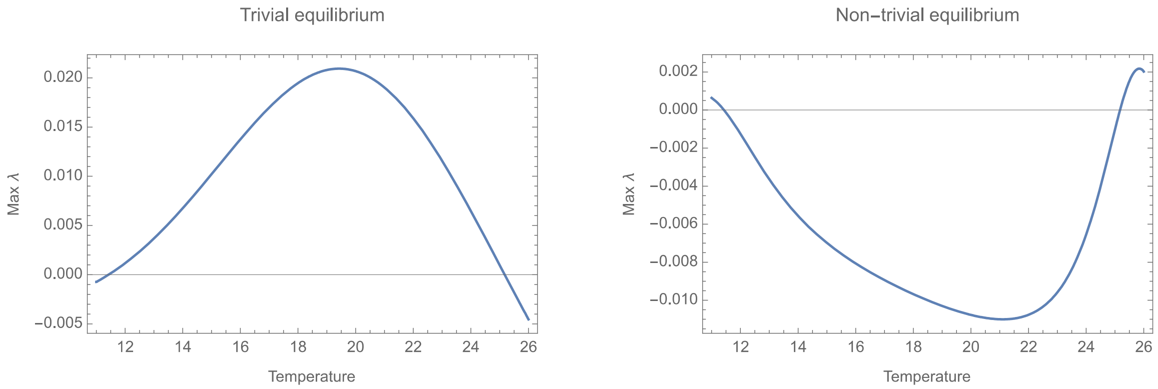


Figure A6. Maximum eigenvalue for A) trivial (axial) equilibrium, and B) non-trivial (interior) equilibrium across temperature.

Dynamics of all state variables

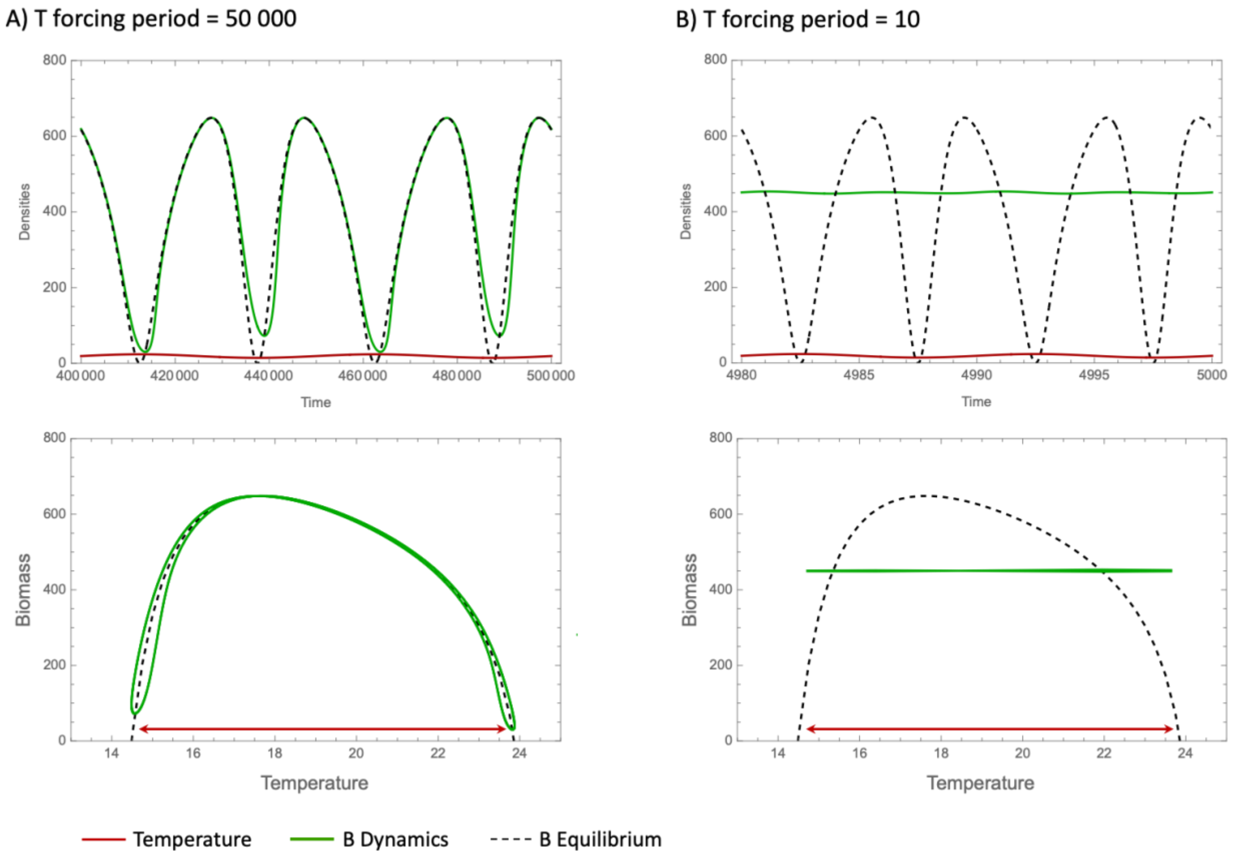


Figure A7. Dynamics of B at additional time scales: 50000 and 10 time units.

For comparison purposes, we show the dynamics of N and Q with temperature again varying between $T_{min}$ and $T_{max}$ (Figure 4 main text). At the time scales displayed here, N is depleted more than would be expected at temperature extremes, with peaks in N corresponding to high and low temperature extremes. With a period of 5000, N displays almost a perfectly sinusoidal (symmetric) pattern, with a period ½ that of the temperature forcing. With faster forcing, it appears that N is more depleted at higher versus lower temperatures, dampening these peaks. The Quota follows expected (equilibrium) values more closely than B and N when the temperature forcing period is 5000, but Q does not reach expected maxima at extreme temperatures, which correspond to higher than expected population biomass (and therefore uptake/assimilation), as well as lower than expected N. Quota is especially lower than expected at temperatures near $T_{min}$. With faster forcing, the Quota levels begin to average out and become more steady.

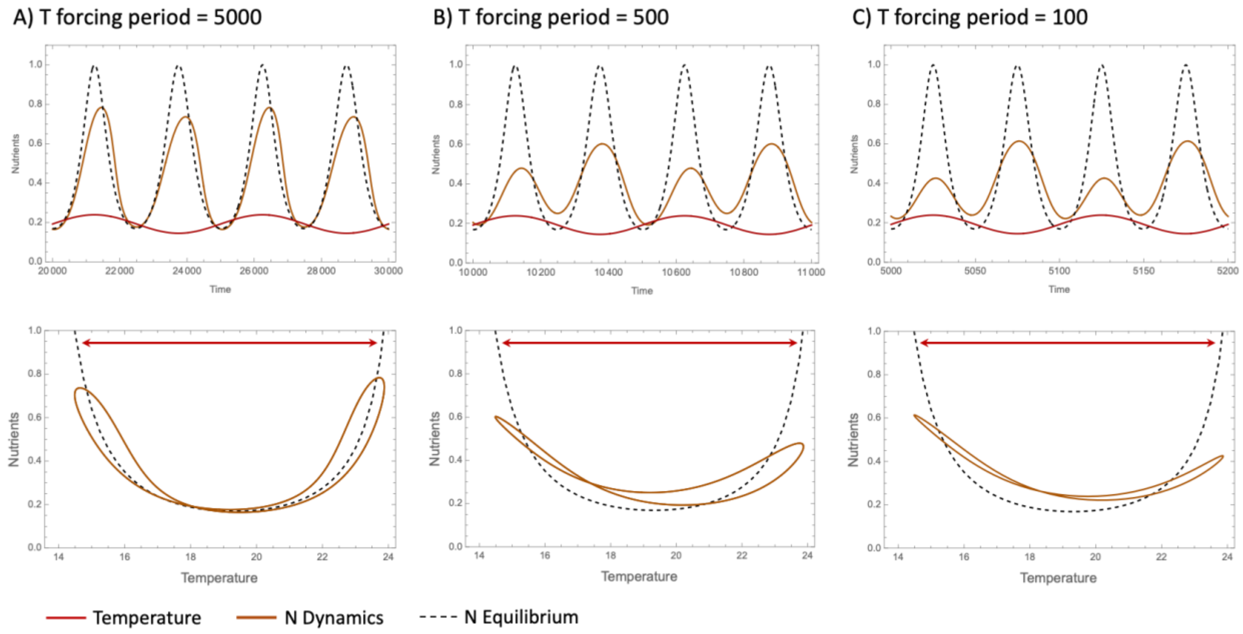


Figure A8. Dynamics of N under sinusoidally varying temperature.

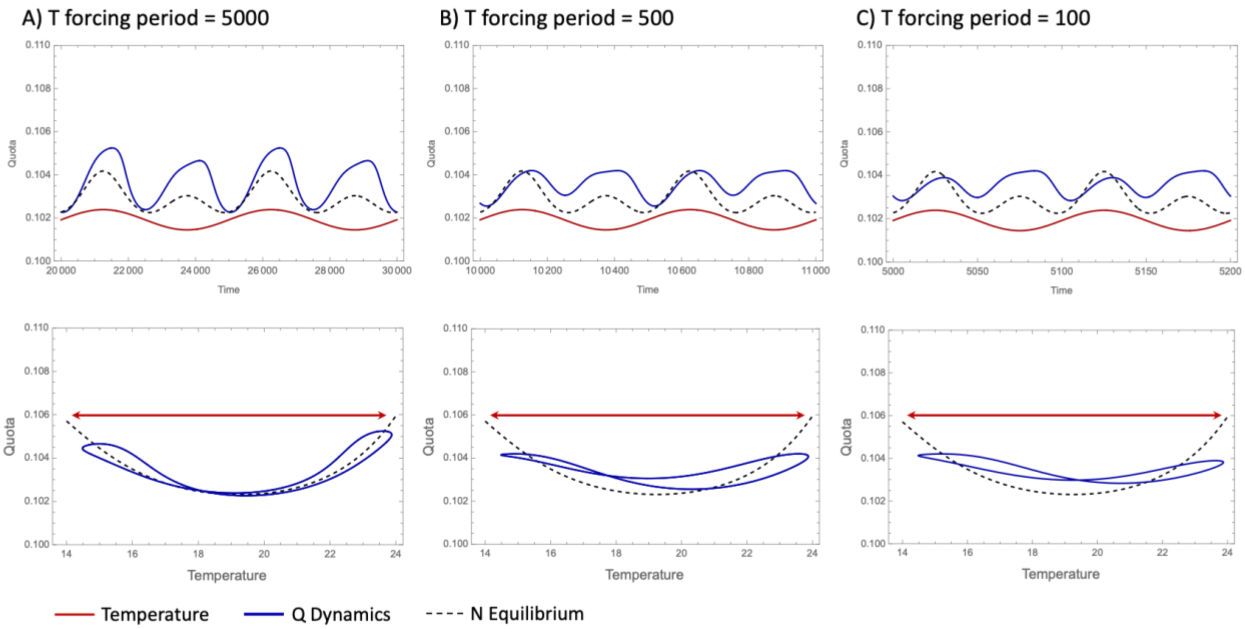


Figure A9. Dynamics of Q under sinusoidally varying temperature.

<u>Dynamics surpassing thermal limits</u>

Temperature regimes that briefly surpass these critical thresholds result in qualitatively similar dynamics as those shown in Figure 4 (main text), except for when forcing is slow enough to allow for population collapse. Even still, collapse occurs more readily at high temperatures than low ones, even though both temperature extremes ought to facilitate population collapse.

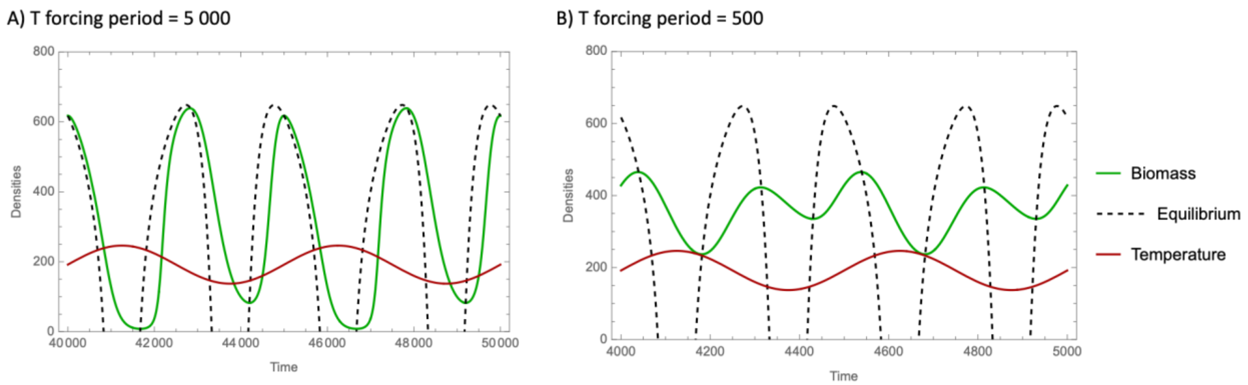


Figure A10. Biomass dynamics with sinusoidally varying temperature temperatures surpass critical thresholds by 0.75 degrees at each extreme (high and low), at two forcing periods: 5000 and 500. Temperature is multiplied by 10 for visualization purposes. Note that slower forcing than this results in collapses at both high and low temperatures to population levels that ought to cause local extinction. Also notice that the dynamics with a forcing period of 500 are qualitatively the same as when temperatures do not exceed the thresholds (see above), with densities just being lower on average. The asymmetric declines in biomass at high versus low temperatures remain clearly visible, however at forcing periods this fast, the population is less influenced by equilibrium dynamics.

<u>Effect of nutrient limitation</u>

Note the following effects of nutrient limitation (increasing $N_0$) on growth and equilibrium temperature responses. First, the entire TPC is diminished such that the thermal niche is decreased, maximum rates of population growth are lower, and the TPC becomes more symmetrical. At the same time, the equilibrium response more closely resembles the TPC in terms of both the curve's symmetry and minimal mismatch. Additionally, since nutrient uptake is limited, there is very little storage in the Quota at all temperatures within the thermal niche (notice Quota invariance in Figure 1). Collectively, this removes asymmetry in the response to high vs. low temperatures and lowers the overall rate of population growth, making the dynamics less responsive to changing temperatures and therefore more invariant.

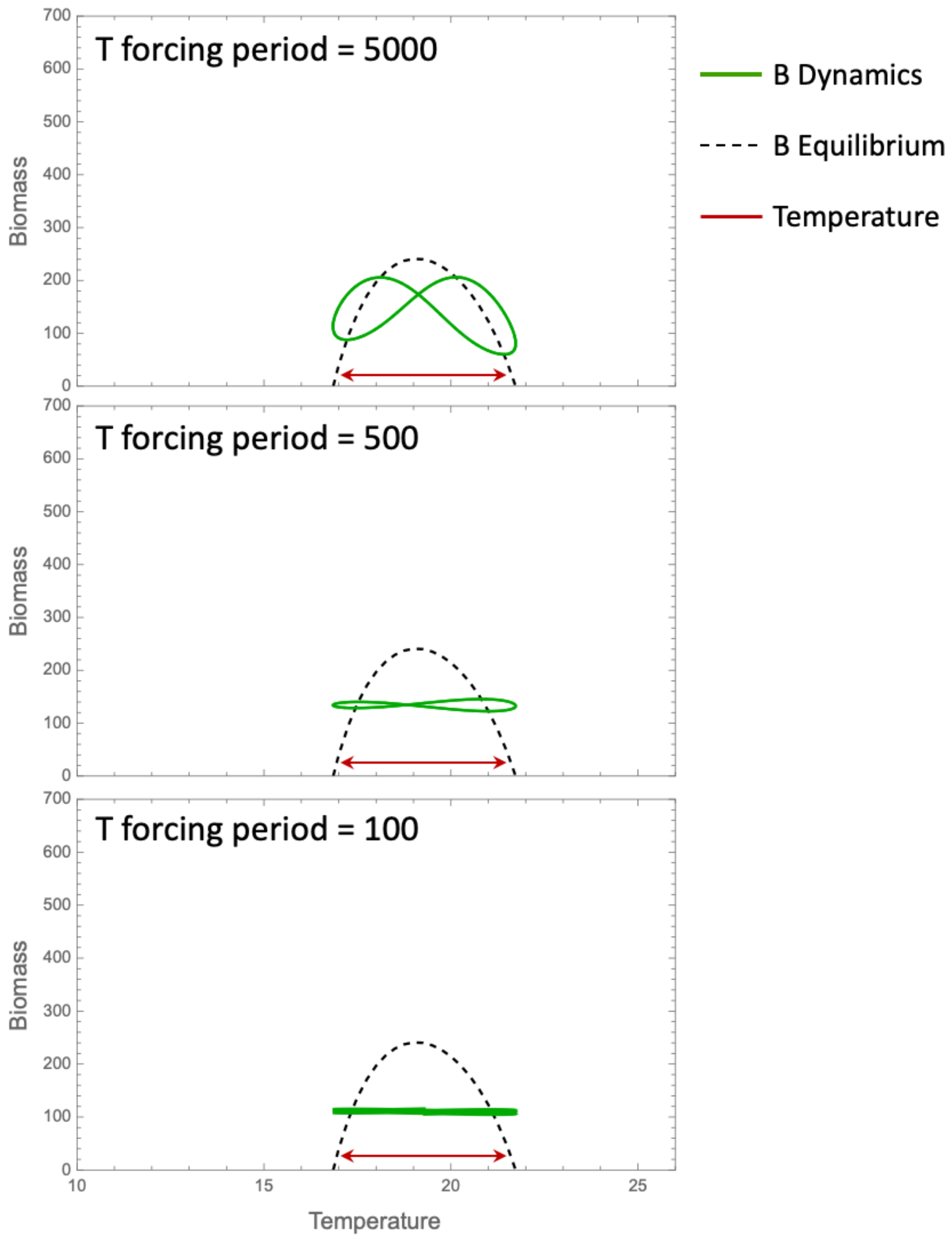


Figure A11. Biomass dynamics with sinusoidal temperature variation across the breadth of the thermal niche. Here, nutrient uptake is limited, with $N_0$ = 2.0.
